# Chromatin as a Coevolutionary Graph: Modeling the Interplay of Replication with Chromatin Dynamics

**DOI:** 10.1101/2025.03.31.646315

**Authors:** Sevastianos Korsak, Krzysztof H. Banecki, Karolina Buka, Abhishek Agarwal, Joanna Borkowska, Haoxi Chai, Piotr J. Górski, Yijun Ruan, Dariusz Plewczynski

## Abstract

Modeling DNA replication requires capturing the interplay of multiple biophysical processes and their effects on chromatin structure. We present RepliSage, a multi-scale framework that integrates three key factors: replication timing, loop extrusion, and compartmentalization. Replication forks act as moving barriers that reorganize chromatin during S phase by interfering with loop extrusion. Our approach combines: (1) fork propagation simulated from single-cell replication timing data, (2) Monte Carlo modeling of loop extrusion and epigenetic state transitions, and (3) 3D reconstruction using OpenMM, where loops are modeled as harmonic bonds and epigenetic states drive compartmental interactions. Chromatin is represented as a graph where both the links and the node states change over time. By adjusting model parameters across cell cycle phases, we reproduce known structural transitions and explore how replication stress alters genome organization. This is the first framework to dynamically integrate all three processes for simulating chromatin architecture during the cell cycle.

## Introduction

This study explores the multi-scale nature of chromatin, a dynamic and hierarchical structure fundamental to genome organization. Chromatin architecture spans from nucleosomes—the basic units of DNA packaging—to higher-order features such as loops and topologically associating domains (TADs). These structures are thought to emerge from the process of loop extrusion, driven by loop extrusion factors (LEFs) and constrained by boundary elements (BEs) (1–3). Beyond these, chromatin exhibits phase separation into A and B compartments, which can further subdivide into subcompartments (4–7), culminating in the spatial arrangement of chromosomal territories (8). The complexity inherent in these scales highlights the challenges of chromatin modeling, both from a physical and biological perspective. A key feature of chromatin, central to this study, is its stochastic nature: chromatin structure is not only multi-scale (9–13) but also dynamic (14–16), with time-dependent processes continuously altering its organization. For instance, the loop extrusion mechanism involves LEFs binding and unbinding along the chromatin (1; 2), forming transient graphs of short- and long-range loops. This process embodies an interplay of order, driven by stabilizing proteins like CTCF, and randomness, arising from the stochastic binding, unbinding, and movement of LEFs. Moreover, compartmentalization has often been associated with the spread of epigenetic marks (17), and is often modeled with Potts-like models (18–24).

DNA replication is a fundamental biological process ensuring genome duplication, occurring during S phase of the cell cycle (25). The eukaryotic cell cycle is divided into four phases: G1, S, G2, and M. Although the actual DNA synthesis is strictly confined to the S phase to prevent genomic instability, the replication process begins earlier with a preparatory step known as origin licensing. This step occurs during late mitosis and early G1 and involves the loading of inactive MCM helicases onto replication origins. Full activation of these origins, and initiation of DNA synthesis, occurs later, specifically during S phase (25; 26). The initiation of replication is temporally regulated in two steps: origin licensing during late mitosis and early G1, followed by origin activation during S phase (26). Only a subset of licensed origins is activated (27; 28), while the remainder are passively replicated by advancing forks (29). Biophysically, replication fork progression is much slower (*u*_rep_ ~ 2 kb/min (30; 31)) compared to loop extrusion speeds (*u*_le_ ~ 0.5 − 2 kb/sec (32; 33)), resulting in a clear separation of timescales between replication and chromatin reorganization. Recent studies indicate that replication forks act as moving barriers to loop extrusion factors (LEFs), weakening TAD borders and loop structures during replication (34; 35), contrasting earlier views where LEFs obstructed forks (36–38). The replication timing program is strongly associated with chromatin compartmentalization: A compartments tend to replicate early in S phase, while B compartments are replicated later (39–43). These compartments reflect underlying epigenetic state, such as histone modifications and DNA methylation, which govern chromatin accessibility and long-range interactions (44; 45). In this context, compartmentalization can be viewed as a spatial manifestation of epigenetic regulation, organizing the genome into transcriptionally active and inactive domains. Furthermore, replication forks have been implicated in the local propagation of epigenetic marks (39; 46; 47), as histone-modifying enzymes and chaperones are recruited to the replisome to ensure faithful transmission of chromatin states during DNA synthesis. Together, these observations suggest that DNA replication is not only temporally aligned with compartmentalization, but may also reinforce epigenetic landscapes that underlie 3D genome architecture.

At the onset of G2/M phase, DNA replication is complete, but sister chromatids remain physically linked, both through cohesion proteins and through unresolved topological entanglements, requiring active mechanisms to separate them in preparation for mitosis (33; 48; 49). Two key processes support this transition: (i) condensin complexes, which extrude chromatin loops more rapidly than cohesins, compacting chromosomes into elongated, supercoiled-like structures essential for mitotic chromosome architecture; and (ii) topoisomerase II (TOP2A) activity, which resolves DNA catenations by transiently cleaving both strands of the DNA helix to allow passage of another DNA segment, ensuring efficient sister chromatid disentanglement (50–52). Together, condensin-mediated loop extrusion and topoisomerase-mediated strand passage orchestrate the dynamic reorganization of replicated chromatin into mechanically stable mitotic chromosomes (48).

Replication stress (RS) is a condition characterized by impaired or stalled DNA replication fork progression, caused by obstacles such as transcriptional interference, nucleotide depletion, or DNA secondary structures (53). To compensate for the reduced fork speed under RS, cells increase origin activation (54). When sustained, RS disrupts the coordination between replication dynamics and chromatin folding, thereby increasing the risk of genome instability, a hallmark of cancer (55; 56). However, previous studies have shown that even under conditions of severe replication stress (57; 58), the global compartmentalization landscape remains largely preserved. This suggests that a primary role of replication forks may be the faithful maintenance of epigenetic organization, and that large-scale perturbations to this structure are relatively resistant to disruption.

In this study, we introduce *RepliSage*, an extension of the LoopSage model (59), designed to simulate DNA replication dynamics integrated with loop extrusion and chromatin compartmentalization. Unlike traditional models focusing only on one-dimensional fork propagation (60–63), RepliSage captures dynamic 3D chromatin alterations by modeling chromatin both as a graph with biophysical attributes and as a molecular dynamics system governed by force fields. It employs a multi-scale stochastic Monte Carlo framework that simultaneously updates chromatin topology and epigenetic states, addressing the complex dynamics of coevolutionary graphs (64–66). RepliSage consists of three components: simulation of replication forks via the *Replikator* module, a stochastic graph-based model incorporating replication and compartmentalization energy terms, and 3D structure reconstruction using OpenMM (Fig.S8). With RepliSage, we simulate the 3D structure of chromatin throughout the cell cycle, encompassing the G1, S, and G2/M phases, and additionally propose a basic model of how replication stress alters chromatin architecture based on the relative dynamics of replication fork progression and origin activation under stress and non-stress conditions. We validated RepliSage using multiple complementary approaches. First, the model accurately reproduces replication timing profiles and Hi-C contact map features, including compartmentalization patterns. We then compared our results to single-cell Hi-C data, confirming that the dynamic behavior of chromatin loops in RepliSage aligns qualitatively with experimental observations. To further assess structural accuracy, we benchmarked RepliSage-generated 3D models against structures produced with NucDynamics, using a set of geometric metrics.

## Results and Methods

RepliSage offers a versatile framework for exploring chromatin organization and DNA replication across the cell cycle. Despite its multi-layered design, the model requires only minimal population-averaged input: replication timing profiles and 3C-based loop data to reconstruct 3D chromatin structures. Its core lies in the optimization of a biophysically-informed energy functional, capturing loop extrusion, replication fork dynamics, and epigenetic interactions. Model outputs are validated against orthogonal datasets, including bulk Hi-C, and single-cell 3D reconstructions from single-cell Hi-C data, demonstrating that RepliSage achieves structural realism grounded in mechanistic assumptions.

### Modelling of Cell Cycle

We simulated the three-dimensional organization of human chromosome 14 throughout the cell cycle, as shown in Fig. 2(a). This analysis was performed for both GM12878 and K562 cell lines (Fig. 2, Fig. S2, and Fig. S3), focusing on a subtelomeric-excluded region of chromosome 14 spanning positions 23,535,000 to 99,174,700 with the cell-cycle specific parameters discussed in Online Methods section. To analyze chromatin folding dynamics, we monitored the evolution of the average loop length over Monte Carlo pseudo-time (Fig. 2(b)). A slight but statistically significant decrease in loop length is observed during the S phase, which we attribute to the dynamic barrier effect imposed by advancing replication forks. In the G2/M phase, condensin activity restricts the formation of long, mobile loops, thereby rapidly guiding the system toward structural equilibrium. To model replication stress conditions, we modified the simulation by increasing the initiation rate function by a factor of four and doubling its variability. Additionally, replication fork propagation speed was reduced by a factor of four. These parameter changes result in fewer than 10% of origins firing under normal conditions, compared to 20–40% under stress conditions. The heightened replication activity under stress leads to a noticeable reduction in average loop size (Fig. 2(b,c)).

**Fig. 1.**
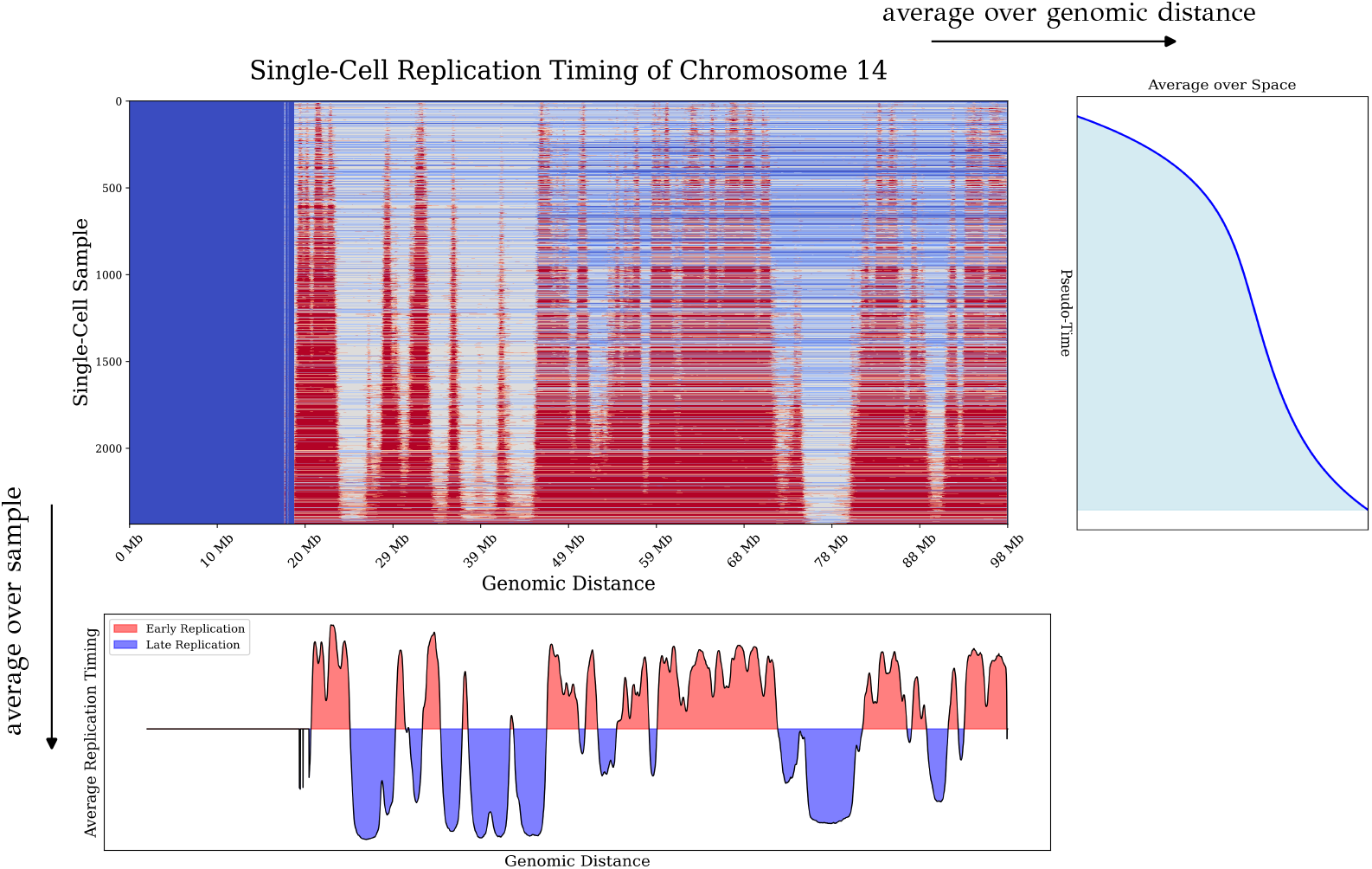
Single-cell replication timing data for chromosome 14. At each genomic locus, replication is represented as a binary state (0 or 1). Temporal averaging across cells yields a replication fraction function *f* (*x, t*), whereas spatial averaging produces a characteristic sigmoid-like transition. Notably, replication timing is highly correlated with chromatin compartmentalization: loci in the A compartment tend to replicate early, while those in the B compartment replicate late. This strong correlation allows compartment identity to be inferred directly from replication timing profiles.

**Fig. 2.**
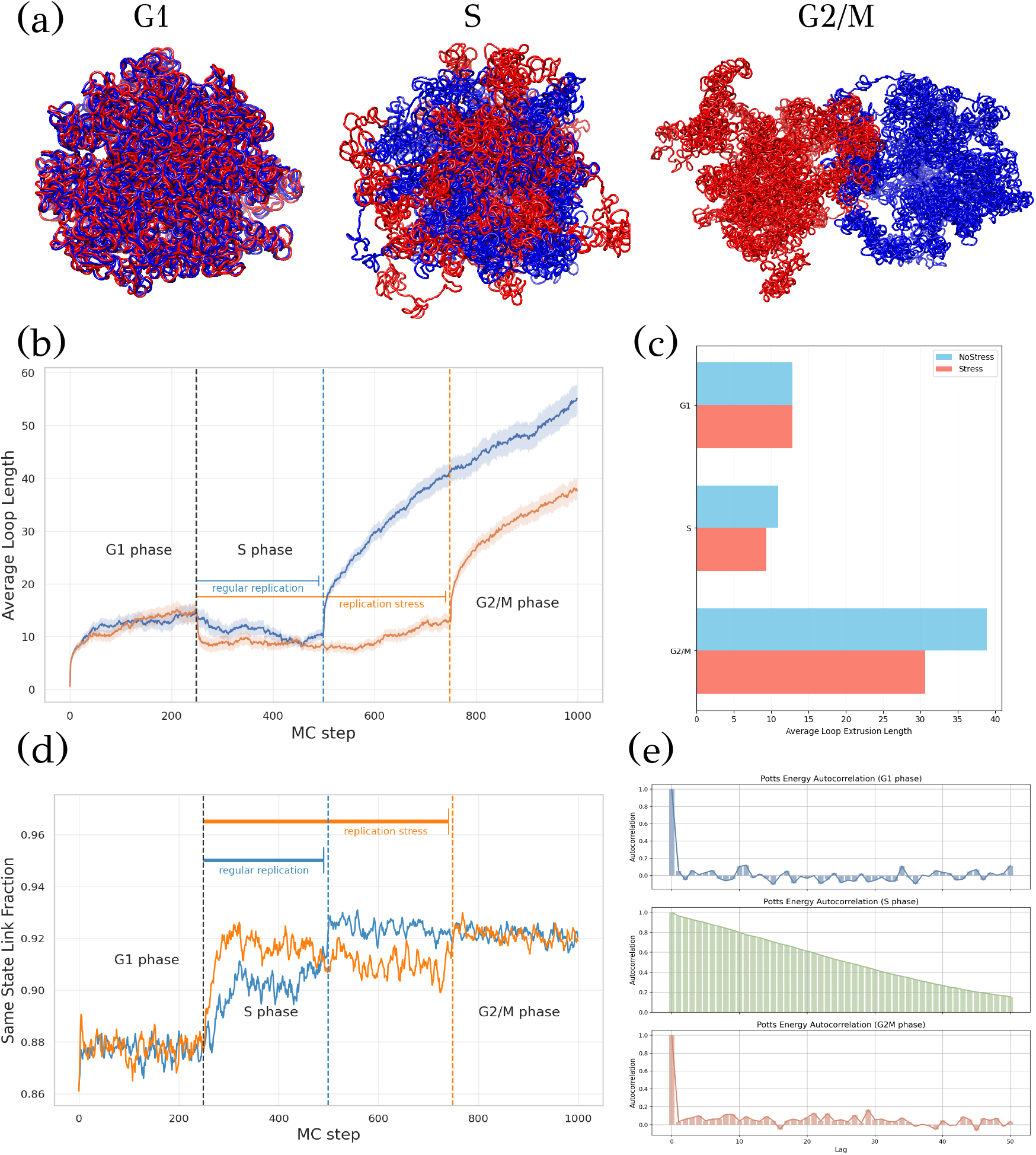
Modeling of chromosome 14 in K562 cells using RepliSage. (a) Representative 3D structures of chromosome 14 during the G1, S, and G2/M phases of the cell cycle. (b) Evolution of the average loop length as a function of Monte Carlo simulation steps, including comparison with replication stress conditions. (c) Average loop size across cell cycle phases under normal and stress-induced replication. (d) Fraction of LEF-mediated links connecting nodes of identical state (i.e., links between nodes with the same sign; for example, nodes with states −2 and −1 are considered equivalent). (e) Autocorrelation in respect of Monte Carlo time of the Potts energy across phases. The system shows no significant autocorrelation in G1 and G2/M phases, while autocorrelation is observed in the S phase due to the out-of-equilibrium dynamics imposed by replication fork barriers.

**Fig. 3.**
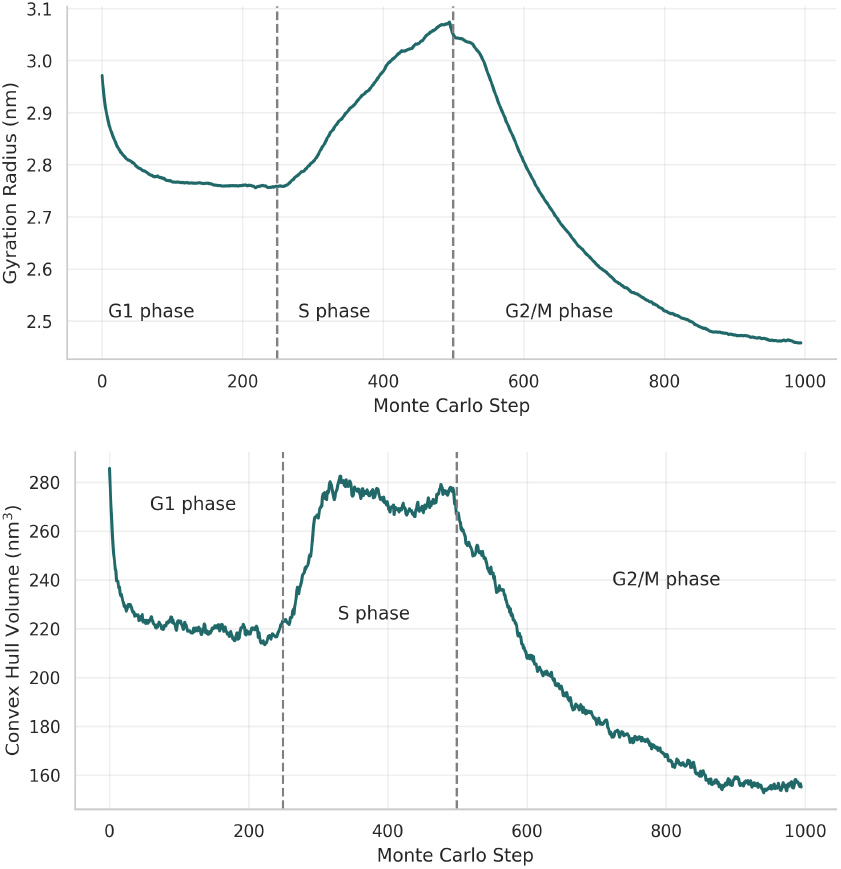
Structural metrics of the 3D chromatin configurations derived from the K562 ChIA-PET dataset. During the G1 phase, the chromatin exhibits compaction. In the S phase, the activity of replication forks leads to a transient decompaction of the structure. In the G2/M phase, increased compaction is observed again, driven by block-copolymer interactions and condensin-mediated binding.

Furthermore, under replication stress conditions, the elevated initiation rate leads to faster synchronization of links connecting loci with identical epigenetic states (Fig. 2(d)). This phenomenon arises because a greater number of replication origins fire earlier, causing replication forks to act as epigenetic organizers that locally promote the alignment of chromatin regions into the A compartment. Notably, despite these differences in synchronization dynamics, the final compartmental organization remains consistent between normal and stress conditions. This result is consistent with experimental findings reporting that large-scale compartmentalization is largely preserved upon different replication stress conditions (57; 58). RepliSage also enables assessment of statistical independence by tracking the autocorrelations of system metrics. As shown in Fig. 2(e), the autocorrelation of the Potts energy confirms that our choice of *f*_mc_ ensures uncorrelated epigenetic states in G1 and G2/M phases. In contrast, higher autocorrelation in the S phase reflects the inherently out-of-equilibrium behavior driven by active replication forks.

In Fig.3 gyration radius and convex hull volume of the single polymer chain are calculated for each structure of the ensemble. These metrics reveal that during the G1 phase, the chromosome is in a compact state. In the S phase, decompaction occurs due to the separation of the two polymer chains and the initiation of replication origins. Subsequently, in the G2/M phase, the chromatin re-compacts, driven by the activity of condensins and enhanced by block copolymer interactions. This dynamic progression illustrates the interplay between replication, structural reorganization, and physical forces shaping chromatin architecture.

### Validation of the Model

Validation of 3D chromatin models is a particularly challenging task. There are two main reasons for this difficulty. First, we lack direct ground truth data describing the true three-dimensional organization of chromatin. Although comparisons with other modeling approaches are possible, such comparisons implicitly assume that at least one of these methods serves as a reliable standard; an assumption that is not necessarily justified. Second, validating against orthogonal experimental datasets, such as those from 3C-type methods, is often problematic. Many studies have shown that direct correlations between simulated all-versus-all distance maps and experimental contact matrices or even between two different experimental heatmaps can be misleading due to biases such as diagonal signal dominance, differences in normalization procedures, smoothing artifacts, or the inclusion of features irrelevant to looping or compartmentalization (67–71). While top-down approaches exist that optimize 3D structures to fit experimental data, they are typically agnostic to underlying biophysical rules and may overfit features that are not biologically meaningful (72–81). Despite these challenges, we believe it is essential to demonstrate that biophysical models maintain some degree of consistency with observed data. For this reason, we undertook a careful and transparent evaluation of RepliSage, aiming to assess its behavior not only through comparison with existing datasets but also by analyzing whether its outputs reflect biologically plausible patterns of chromatin organization.

RepliSage integrates biophysical modeling across multiple chromatin scales. At the TAD level, it incorporates a modified version of LoopSage (59), which we have previously validated by comparing loop-specific features between simulations and experimental data. While RepliSage centers on chromosome-scale modeling rather than loop extrusion itself, faithfully representing loop extrusion is essential because it directly interacts with replication processes and the transmission of epigenetic states. At the compartment scale, RepliSage employs an epigenetic Potts field derived from replication timing data. Since replication timing strongly correlates with chromatin compartmentalization, the resulting epigenetic state patterns closely mirror Hi-C derived compartments. As shown in Fig. 4(a–b), the ensemble-averaged epigenetic states achieve over 80% Pearson correlation with Hi-C compartment profiles. To further validate large-scale organization, we compared the first eigenvectors of inverse distance matrices from simulated ensembles (Fig. 4(c)) with those from Hi-C data (Fig. 4(d)), observing correlations above 60%. In contrast, random walk ensembles, lacking compartmental structure, show near-zero correlation, supporting the biological relevance of RepliSage output. Although we computed direct correlations between model-derived and experimental heatmaps, we excluded them from the main analysis due to their susceptibility to technical artifacts.

**Fig. 4.**
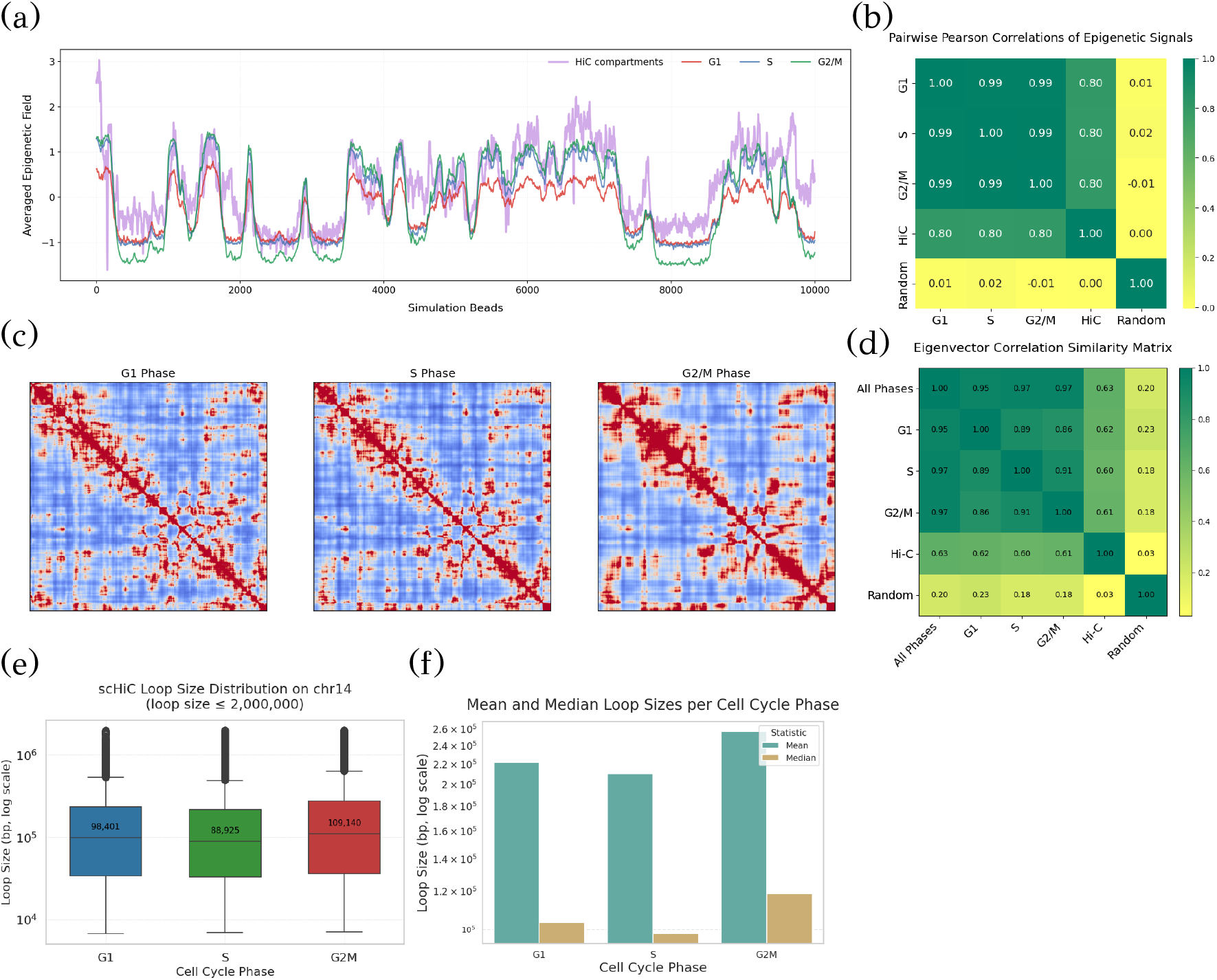
Validation of the RepliSage model. (a) Ensemble-averaged epigenetic field compared to Hi-C compartmentalization profiles across different cell cycle phases. (b) Pearson correlation between simulated and experimental compartment signals. (c) Averaged inverse all-versus-all distance heatmaps from RepliSage across cell cycle phases. (d) Correlation between the first eigenvector of simulated distance maps and Hi-C compartment eigenvectors. (e) Distribution of interaction distances derived from single-cell Hi-C data. (f) Phase-specific average and median interaction lengths from single-cell data.

Since RepliSage is primarily designed to model DNA replication dynamics, we performed qualitative validation using single-cell Hi-C datasets. A key reference was the dataset from Chai et al. (82), which provides insight into the distribution of loop sizes across different cell cycle phases. To analyze these patterns, we generated phase-specific datasets by merging overlapping anchors and retaining statistically significant interactions supported by at least three contacts. Interactions exceeding 2000 kb were excluded, as they are unlikely to result from loop extrusion. This analysis revealed a consistent trend: in the S phase, average loop lengths are shorter than in G1, while in G2/M, loops tend to be longer, likely due to condensin-mediated compaction (Fig. 4(e–f)).

### Low resolution models of chromosomes

To gain deeper insights into chromatin structural dynamics throughout the cell cycle, we generated low-resolution (100 kb) 3D models for the K562 (Fig. 5) and GM12878 (Fig. S9) cell lines. Violin plots (Fig. 5(a)) confirm that S phase exhibits the shortest loop sizes. Larger chromosomes showed longer loops, possibly due to low resolution in smaller chromosomes. The results were consistent with previous high-resolution analyses, showing a robust decrease in average loop size during S phase under replication stress compared to normal conditions (Fig. 5(b)), likely due to increased replication fork activity acting as moving barriers to loop-extruding factors (LEFs).

**Fig. 5.**
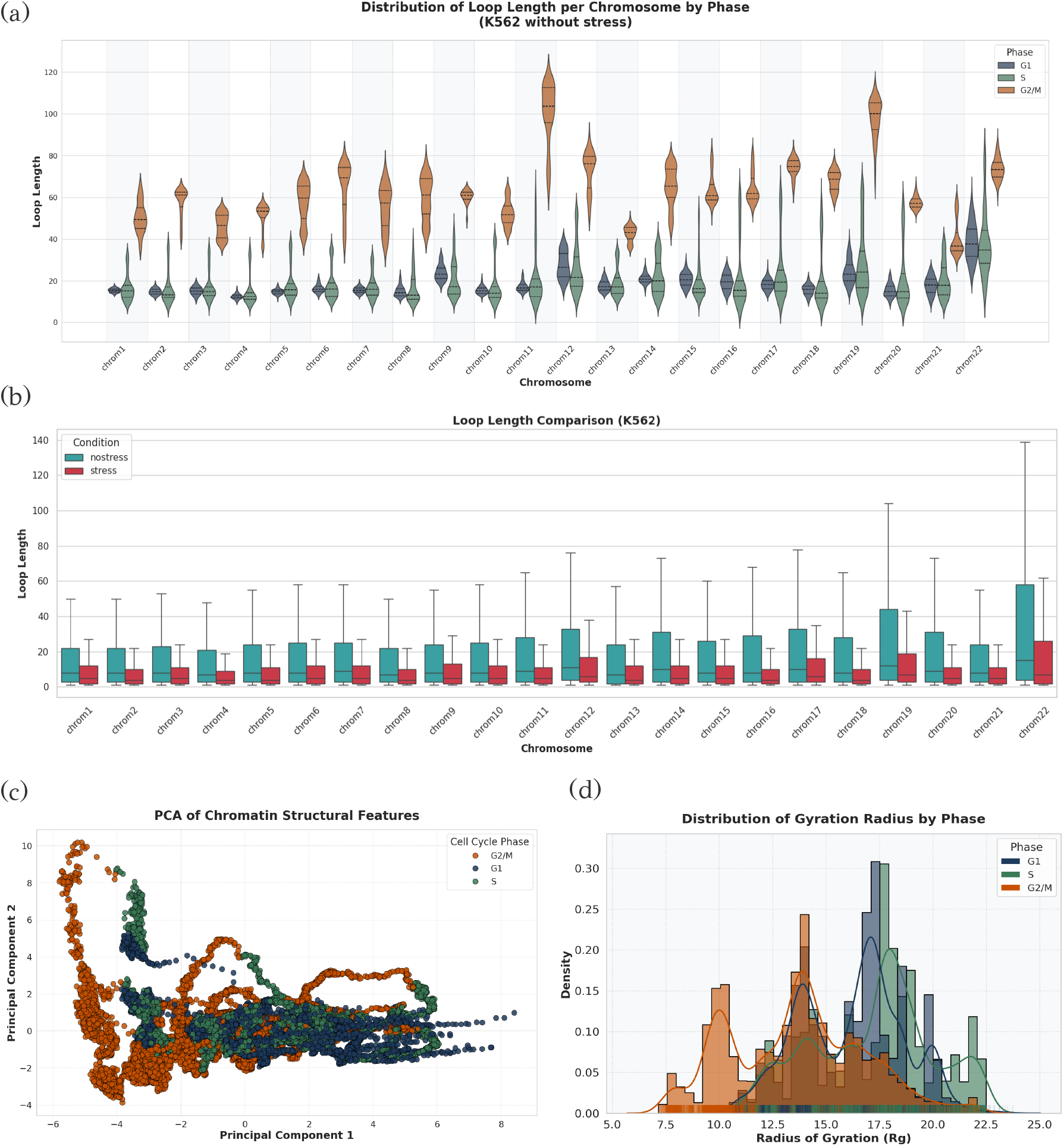
Chromosome-scale structural changes across replication and cell cycle phases as predicted by RepliSage using CTCF ChIA-PET input data from the K562 cell line. (a) Distribution of loop lengths across distinct cell cycle phases. During the S phase, loops tend to be significantly shorter, consistent with the barrier effect exerted by active replication forks. In contrast, the G2/M phase is characterized by longer loop structures, likely reflecting condensin-mediated chromatin compaction. (b) Comparison of loop length distributions under normal and replication stress conditions. Replication stress during the S phase results in a marked reduction of average loop size, indicating disrupted loop extrusion dynamics. (c) Principal Component Analysis (PCA) was applied to structural descriptors (as defined in the *Definition of Structural Metrics* section) computed for each individual polymer conformation within the ensemble of each chromosome. Each conformation was encoded as a high-dimensional feature vector capturing its geometric attributes. The dimensionality reduction was carried out in an unsupervised manner, without incorporating cell cycle phase information. The resulting two-dimensional projection exhibits weak clustering by cell cycle phase, suggesting the presence of subtle, phase-specific structural signatures. Each point in the plot corresponds to a single 3D structure, embedded in the PCA-derived 2D feature space. (d) Distribution of the radius of gyration *R*_*g*_ across all chromosomes and cell cycle phases. The histogram indicates global chromatin decompaction during the S phase, as evidenced by a shift toward higher *R*_*g*_ values. Conversely, the G2/M phase shows increased compaction, reflected by a shift toward lower *R*_*g*_ values. These observations support phase-dependent modulation of chromatin folding.

To quantitatively assess the structural properties of chromatin across the cell cycle, we computed a range of geometric descriptors, as detailed in the Online Methods (see ‘Definition of Structural Metrics’ section). These descriptors were used to perform principal component analysis (PCA), as shown in Fig. 5(c). The PCA results revealed clear clustering of chromatin conformations according to cell cycle phase, with each phase forming distinct groups in the reduced-dimensional space. Furthermore, analysis of the radius of gyration *R*_*g*_ across chromosomes (Fig. 5(d)) demonstrated a consistent trend of chromatin decompaction during the S phase, followed by increased compaction during the G2/M phase relative to G1. These findings underscore the phase-specific reorganization of chromatin structure during cell cycle progression.

### Comparison with Models from Single-Cell Data

To evaluate whether our results are consistent with previous studies on 3D chromatin dynamics during cell cycle progression, we utilized the recent dataset from Chai et al. (2025) (82), focusing on the K562 cell line. In that study, cells were classified into G1, S, and G2/M phases based on the expression of phase-specific genes. Principal component analysis (PCA) was performed, and the contact data were projected onto the first two principal components to group cells into pseudo-bulk metacells, representing distinct stages along the cell cycle pseudo-time. Following their methodology, we employed the NucDynamics software (83) to reconstruct 3D chromatin structures for each metacell. This allowed us to investigate the overall patterns of chromatin structural changes and compaction across the cell cycle, which we then compared to analogous results generated using RepliSage for the same K562 cell line.

To quantitatively assess chromatin compaction throughout the cell cycle, we calculated the Radius of Gyration (Rg) and the convex hull volume for each chromosomal structure. The results are presented in Supplementary Figures S4, S5 and S7. In the NucDynamics-based analysis, both Rg and convex hull volume gradually increased from G1 to S phase, indicating a relaxation of chromatin, and then decreased upon entry into the G2/M phase, suggesting a re-compaction of chromatin. A comparable pattern was observed with the RepliSage results (see Figure 5 (d)), thereby supporting the validity of our approach. Notably, although RepliSage does not incorporate contact data in its modeling, the replication dynamics it infers align closely with known chromatin compaction trends from other studies (34) as well as with our own 3D structural reconstructions. This consistency highlights the robustness of RepliSage in capturing broad chromatin behavior across the cell cycle, despite relying on an orthogonal data modality.

### Parameter Study

As detailed in the *Online Methods, RepliSage* exhibits strong sensitivity to its parameter settings. Therefore, gaining at least a qualitative understanding of its behavior under perturbations of key parameters is essential. Among these, one of the most critical is the biological temperature, or order parameter, denoted as *T*_mc_. To facilitate a systematic parameter study, we focused our analysis on a representative subregion of chromosome 14, specifically the genomic interval [70,835,000, 98,674,700]. Within this region, each individual polymer conformation was modeled as a chain of 2,000 beads. This reduction in genomic span was chosen to enable faster simulations across different parameter sets. Figure 6(a) shows how the average loop length varies as a function of the order parameter *T*_mc_. Lower values of *T*_mc_ lead to longer, more stable loops, while higher values result in shorter loops. This behavior can be interpreted by viewing *T*_mc_ as analogous to a biological temperature: higher values introduce greater kinetic noise, increasing the frequency of LEF unbinding and reducing the time available for loop extrusion. Conversely, lower values lead to more stable configurations in which LEFs remain bound longer, allowing loops to grow and stack. To observe clear differences in loop size across cell cycle phases, a lower *T*_mc_ is preferable, as it favors energy-minimizing moves over stochastic transitions. Under these conditions, the effect of replication forks acting as dynamic barriers, most pronounced in the S phase, is more clearly captured. The behavior of the average epigenetic field, shown in Fig. 6(b), is consistent with classical results from Ising and Potts models. At high temperatures, the epigenetic states fluctuate randomly, producing biologically uninformative configurations. At lower temperatures, state transitions are suppressed, leading to more ordered configurations and more negative average field values. The presence of replication forks effectively shifts the Potts energy minimum, reinforcing the organization of epigenetic domains.

**Fig. 6.**
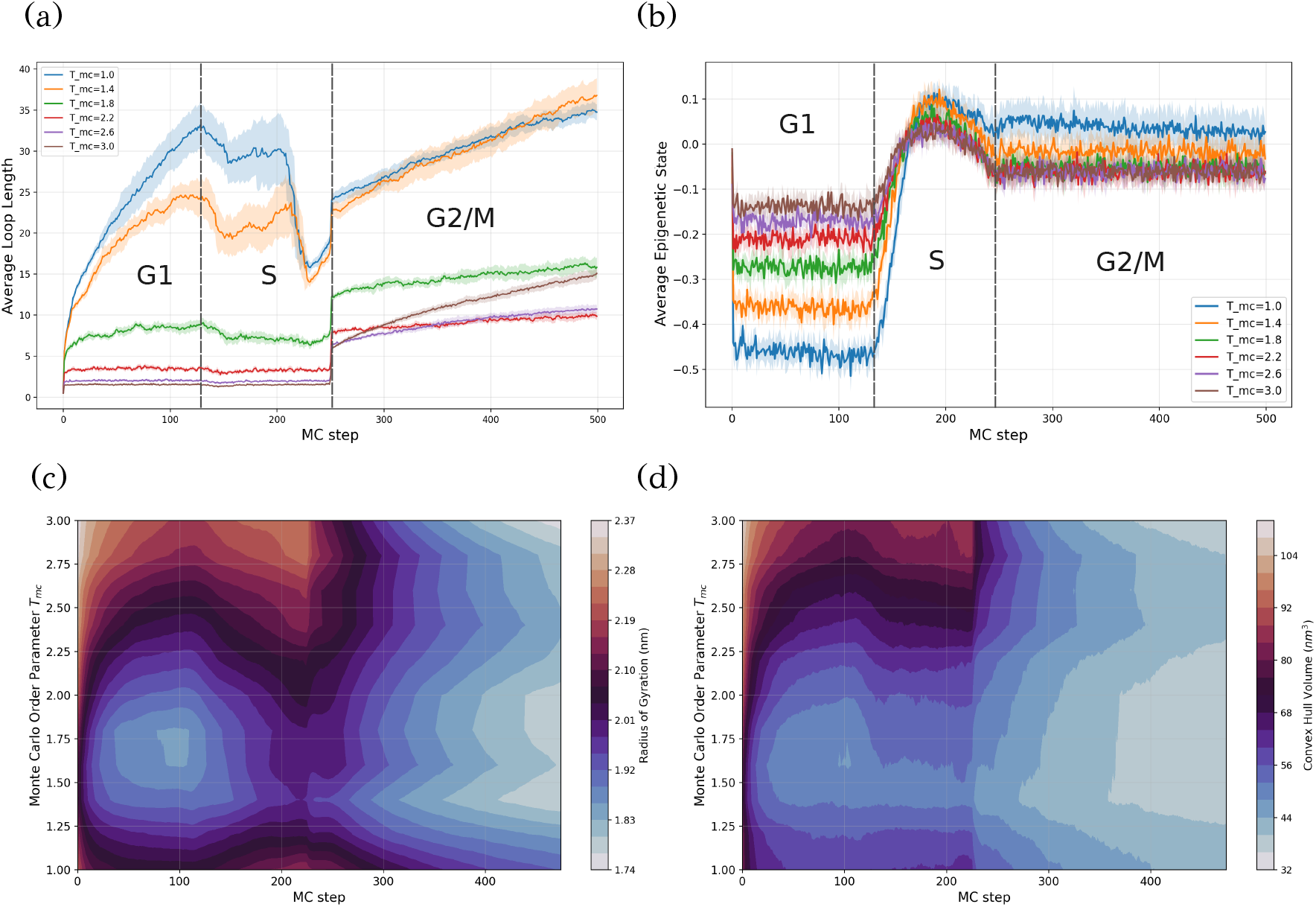
Parameter study of RepliSage. Results are based on 10 independent simulations using CTCF ChIA-PET data from the GM12878 cell line, focused on the illustrative region [70835000, 98674700] of chromosome 14. For each value of *T*_mc_, 10 independent runs of the RepliSage algorithm were performed and subsequently averaged. (a) Average loop length as a function of the order parameter *T*_mc_. Elevated values of *T*_mc_ correspond to shorter loops, whereas lower values reproduce previously reported trends: moderate compaction during S phase followed by enhanced compaction in G2/M phase. (b) Average epigenetic state, quantified as the mean value ⟨*s*⟩ across all nodes in the chromatin graph, plotted as a function of *T*_mc_. A discrete jump in ⟨*s*⟩ reflects replication-induced epigenetic reconfiguration driven by replication fork progression. (c) Contour plot depicting the evolution of the radius of gyration as a function of both *T*_mc_ and Monte Carlo time. A range of critical *T*_mc_ values is observed at which chromatin compaction is maximized. (d) Contour plot of the convex hull volume as a function of *T*_mc_ and Monte Carlo time.

The applied Potts model includes, among others, two key coefficients: one governing the strength of interactions between chromatin segments, and the other representing the influence of the epigenetic field (see Eq. 10). An interesting question arises regarding the role of these two coefficients (Fig. S10); specifically, whether changes in the balance between the interaction and field terms can drive large-scale reorganization of epigenetic patterns. In our simulations, although the global epigenetic landscape remains largely preserved across different parameter regimes, we observe local rearrangements and highly correlated behavior when the interaction term dominates over the field term. This behavior is consistent with known results from statistical physics, where phase transitions, interpreted in the biological context as large-scale epigenetic reprogramming or decoupling from initial chromatin states, emerge in systems where long-range interactions prevail (84–88). In our chromatin model, where interaction matrices are defined solely by extrusion-mediated loops, only short-range correlations are present. Consequently, the system remains in a robust phase below the transition threshold, consistent with experimental findings that chromatin compartments are stably maintained across all phases of the cell cycle and under both stress and non-stress conditions (57; 58). According to our model, two conditions would be required for loop extrusion alone to drive compartmental organization: (1) that loop extrusion itself plays a central role in establishing compartments, and (2) that it generates long-range loops spanning compartment-scale distances. However, current evidence suggests that neither condition holds biologically. Therefore, loop extrusion is not only insufficient to generate compartmentalization (89; 90), but its local and short-ranged nature (33; 72; 91) is precisely what explains the remarkable stability of the compartmental pattern, even under significant DNA stress.

An important and non-trivial observation concerns how the three-dimensional chromatin structure is influenced by variations in the Monte Carlo parameters. This connection is far from obvious, as the internal graph states of the model, defined in one dimension, just ultimately give rise to a spatial structure in three dimensions, where the system has significantly more degrees of freedom. Figures 6(c) and 6(d) present contour plots of the radius of gyration and convex hull volume, respectively, as functions of both *T*_mc_ and Monte Carlo time. As expected from prior results, the chromatin structure is more compact in the G1 and G2/M phases. However, a particularly interesting result is the emergence of a critical temperature region around 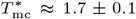, where compaction reaches a maximum. This optimal folding does not occur at the lowest temperatures, but rather within an intermediate range, suggesting that a certain level of stochasticity is necessary to achieve biologically realistic 3D organization. Notably, this behavior is consistent across all cell cycle phases, indicating a robust temperature-dependent folding regime within the model.

## Discussion

In this study, we introduced a novel computational framework to model chromatin as a coevolutionary graph, in which both the epigenetic states of chromatin segments (nodes) and the physical connections between them (links) evolve over time. This dual-level dynamical system captures the interplay between chromatin structure and epigenetic regulation, particularly during the complex biological process of DNA replication.

DNA replication has long attracted interest from physicists aiming to describe it through analytical models and stochastic frameworks (60–63; 92–95). Early models typically relied on simplifying assumptions, such as uniform fork speeds and homogeneous firing rates, to derive transport equations that describe the temporal and spatial dynamics of replication forks. More recent stochastic approaches have incorporated spatial inhomogeneities and probabilistic origin activation, but largely remain one-dimensional in nature (60; 61). Although replication kinetics have been extensively studied theoretically, the link between one-dimensional replication fork progression and three-dimensional chromatin architecture remains insufficiently understood. Only recently have studies begun to explore this relationship (96–99). While these efforts represent important progress, they often overlook key aspects such as the role of SMC proteins, epigenetic mark propagation, and chromatin reorganization during the G2/M phase. By treating the cell cycle as a phase-transition system, RepliSage dynamically switches simulation parameters to reflect transitions between G1, S, and G2/M phases. Our approach establishes a direct link between 1D chromatin dynamics, where replication and loop extrusion are modeled probabilistically, and 3D structure, simulated through OpenMM-based molecular dynamics. This dual-level representation enables the generation of realistic 3D chromatin conformations while supporting systematic parameter studies that reveal how structural metrics evolve under different biological conditions, including replication stress.

Moreover, the resulting predictions can be compared with experimental datasets, including bulk Hi-C and single-cell chromatin conformation data (71; 100), providing a powerful platform for hypothesis testing and model validation. Taken together, RepliSage represents a minimal yet comprehensive modeling framework that enables a new class of biologically grounded simulations, bridging the gap between dynamic replication processes and their structural manifestations in the nucleus. Furthermore, our model has been validated through multiple complementary approaches. These include testing various biophysical assumptions by integrating them into the framework, comparing results against orthogonal experimental datasets, and evaluating consistency with independent single-cell models. This highlights that chromatin model validation is not a one-dimensional process, as it requires a careful interpretation of experimental data, followed by its translation into mathematical and physical terms. For instan
ce, we compare the contact heatmaps generated by *RepliSage* with orthogonal Hi-C data, as well as with heatmaps produced by random walk models. These comparisons consistently show stronger correlations with our non-random, physically grounded model. Additionally, we demonstrate that the average epigenetic state predicted by our Potts model aligns well with the first eigenvector of the Hi-C contact matrix, indicating that replication timing curves alone are sufficient to reconstruct key structural features observed in experimental data. One of the most significant validations of our approach comes from its comparison with single-cell Hi-C data. Remarkably, *RepliSage*, using only bulk replication timing as input, can reproduce structural patterns that closely resemble those found in single-cell datasets. This result provides mutual validation of both the single-cell data and our modeling framework.

In summary, *RepliSage* offers a robust and extensible framework for modeling chromatin dynamics during DNA replication. It captures key biophysical features, such as loop extrusion and replication-driven remodeling, while requiring only minimal assumptions and no reliance on extensive single-cell data. Despite its simplicity, the model achieves strong predictive power and can be easily extended to incorporate additional mechanisms, such as chromatin knotting or the impact of double-strand breaks. This makes it a versatile tool for current and future studies in genome biology.

## Data Availability

RepliSage model can be found as an open-source project in https://github.com/SFGLab/RepliSage. The updated version of LoopSage can be found in https://github.com/SFGLab/pyLoopSage or via PyPI in https://pypi.org/project/pyLoopSage/.

## Acknowledgments

This work has been supported by Polish National Science Centre (2020/37/B/NZ2/03757). Research was co-funded by Warsaw University of Technology within the Excellence Initiative: Research University (IDUB) programme. Computations were performed thanks to the Laboratory of Bioinformatics and Computational Genomics, Faculty of Mathematics and Information Science, Warsaw University of Technology using Artificial Intelligence HPC platform financed by Polish Ministry of Science and Higher Education (decision no. 7054/IA/SP/2020 of 2020-08-28). The work was co-supported by National Institute of Health USA 4DNucleome grant 1U54DK107967-01 and “Nucleome Positioning System for Spatiotemporal Genome Organization and Regulation”. Finally, we also gratefully acknowledge the Koren Lab for providing access to single-cell replication timing data whenever needed.

## Online Methods

This section provides additional details about the experimental procedures and a more in-depth explanation of the RepliSage model. It is intended for advanced readers seeking a deeper understanding of the methodology, parameterization, and assumptions underlying our approach. We strongly encourage users who wish to apply RepliSage in their own research to consult this section carefully. A thorough understanding of the model parameters, their biological relevance, and their impact on simulation outcomes is essential for proper usage and meaningful interpretation of the results.

### Input data

RepliSage operates on two primary input types: (i) *replication timing profiles*, and (ii) *chromatin loop data* from 3C-type experiments to define CTCF-mediated interactions. By default, RepliSage internally loads single-cell replication timing data from the Koren lab (101), removing the need for users to provide this input. However, alternative datasets can be easily substituted if desired. In this study, we used CTCF ChIA-PET data from ENCODE for two cell lines: GM12878 (ENCSR184YZV) and K562 (ENCSR597AKG). Our primary analyses focus on K562, as single-cell replication data is available for this line, enabling more robust validation. Moreover, because K562 is a cancer cell line, it more realistically reflects replication stress conditions - one of the central use cases RepliSage was designed to model.

For validation, we compared RepliSage outputs with orthogonal Hi-C datasets (ENCSR968KAY for GM12878 and ENCSR545YBD for K562), and qualitatively assessed key biophysical features such as loop sizes and chromatin compaction using single-cell benchmarks (82). To further investigate replication dynamics under stress, we also incorporated Hi-ChIP data, enabling direct comparison of loop size distributions between normal and stress-induced replication conditions. These Hi-ChIP datasets are unique to this study, generated in our laboratory, and are provided in the Supplementary Materials. Additionally, we assessed the consistency of RepliSage predictions by comparing them with single-cell-based chromatin structure models obtained using the NucDynamics framework. Although RepliSage operates independently, the strong agreement between the two approaches underscores its capacity to reproduce features characteristic of single-cell-level chromatin organization.

### ChAIR

The ChAIR method was performed as described in Chai et al. (82). Briefly, crosslinked cells were treated with restriction enzyme digestion, proximity ligation, and transposition *in situ*. The intact cells were then processed using the 10x Genomics Multiome platform following the manufacturer’s instructions.

### Cell Cycle Phasing by Gene Expression

Cell cycle scores were assigned to each cell and projected onto the PCA plot, following the methodology outlined in the Seurat cell cycle vignette^1^. We then identified the centroids for each cell-cycle phase (G1, S, and G2/M) in PCA space using the first two principal components. For our calculations, we defined the centroid of a given cell-cycle phase as:

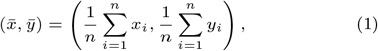

where *x*_*i*_ and *y*_*i*_ represent the PC1 and PC2 coordinates of cell *i*, and *n* is the number of cells in the specific cell-cycle phase.

We then computed the Euclidean distance from a given cell to the centroids of the previous and later cell-cycle phases to define the differential distance (DD):

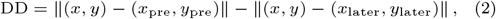

where (*x, y*) are the coordinates of the current cell, and (*x*_pre_, *y*_pre_) and (*x*_later_, *y*_later_) denote the centroids of the previous and later phases, respectively.

Cells were subsequently ranked according to their DD values in descending order within each phase. Cells with higher DD values are closer to the previous phase, while those with lower values are nearer to the later phase, indicating their progression through the cell cycle.

For metacell analysis, we aggregated single cells from the same phase, starting from the lowest to the highest DD rank, to form metacells, each comprising approximately 1 million chromatin contacts.

### NucDynamics simulations

Single cell NucDynamics simulations on the metacell data were performed with the default parameters. That is, for each metacell hierarchical simulation scheme was employed for a single conformational model with resolutions: 8Mb, 4Mb, 2Mb, 400kb, 200kb and 100kb. The output of the simulations were .pdb files with 3D coordinates used subsequently for the radius of gyration and convex hull volume calculations.

### RepliSage Model Structure

The RepliSage method consists of three distinct simulation modules: (1) a replication simulator (*Replikator*) that models stochastic origin firing and the propagation of replication forks; (2) a stochastic simulation of loop extrusion, capturing its dynamic interplay with replication forks and chromatin compartmentalization; (3) a physical modeling component based on OpenMM (102; 103), which reconstructs the 3D chromatin structure using a specified force field. RepliSage supports simulation of different cell cycle phases through adjustable parameters. All components are fully reproducible and publicly available via GitHub and PyPI. For ease of use with CUDA, a Docker container is also provided, enabling third parties to deploy the model with minimal effort.

### Data Preprocessing

Before running the replication simulation in RepliSage, it is essential to perform a preprocessing step to extract the necessary parameters. We define the initiation rate *I*(*x, t*) as the probability per unit time and per unit length that a replication origin at position *x* initiates replication at time *t*. Given this initiation landscape *I*(*x, t*), together with the replication fork velocities *v*_*k*_, the replication process can be simulated using a straightforward Monte Carlo algorithm. Following an approach similar to (62), we first compute the replication fraction *f* (*x, t*) by averaging single-cell replication timing data across time (Fig. 1(a)). We then assume that the initiation rate at each genomic position follows a Gaussian profile in time:

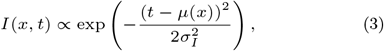

where *µ*(*x*) denotes the mean replication time at position *x*, obtained directly from the average replication timing curve. The variance *σ*_*I*_ is assumed constant across the genome and is set proportionally to the total replication time *T*_*r*_. In our replication model, *T*_*r*_ is provided as an input parameter. Consequently, the progression of replication is primarily controlled by the fork speed and the number of active replication forks. It is important to note that the initiation probabilities *I*(*x, t*) are not sampled directly from the Gaussian distribution (Fig. S1(a)). Instead, replication origins are stochastically tested for firing at discrete time steps throughout the simulation. To ensure that this iterative sampling procedure yields a firing time distribution consistent with Equation (3), the values of *I*(*x, t*) were renormalized accordingly.

The task of accurately estimating the parameters of the normal distribution only from the binary single cell states is extremely challenging. Therefore, our approach is very similar to the one proposed by Yousefi et al (62). Consequently, we assume that our model is good enough as long as averaged independent experiments of the replication simulation, which we will describe in the following section, can reconstruct indeed the averaged replication timing curve.

According to the paper (62) we can calculate the distribution of speeds directly from the slopes of the replication curve. In this case, we assume that the replication fork speed-meaning both sides of the fork move at the same rate-follows a normal distribution with mean *µ*_*v*_ and standard deviation *σ*_*v*_:

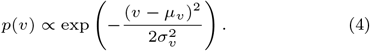

The parameters *µ*_*v*_ and *σ*_*v*_ are determined as the mean and standard deviation of the slopes between consecutive optima of the replication curve *f*. Despite the fact that our pre-processing approach is relatively simple, we were able to reconstruct the average replication curve only with these assumptions (Fig. S1(b)).

### Replication Simulation (Replikator)

Initially, the initiation rate and replication fork speed are imported into the replication simulation. Replikator is a simplified model designed to simulate the stochastic propagation of replication forks (Fig.S1(d)). The primary assumption of this model is the stochastic nature of origin firing. In a single run of Replikator, only a subset of origins initiates replication, while other origins are passively replicated. Despite this simplification, our objective is to ensure that the model accurately reconstructs the average replication timing curves used as input in RepliSage. Consequently, we have designed Replikator to produce highly correlated average replication fork trajectories that align with experimental data (Fig.S1(b)).

In each epoch of the simulation, we evaluate all monomers by drawing a random number in the interval [0, 1] for each monomer. If the number is less than the initiation rate *I*(*x, t*) for the given time *t*, the Monte Carlo move is accepted, and the origin fires. Upon origin firing, we draw a velocity from the probability distribution *p*(*v*) defined in Equation (4). In the second phase of the epoch, we propagate each replication fork a distance *r* = *v*(*t* − *t*_0_), where *t*_0_ is the initiation time of a specific origin. The simulation terminates when the entire DNA is replicated. The trajectories of replication forks and the simulation-estimated replication fraction function are obtained as outputs of the simulation.

### Stochastic Model

The stochastic model’s energy is composed by three factors,

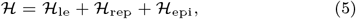

where each one of them aims to model different aspects of biophysics of chromatin. In this sense, ℋ_*le*_ factor is split into folding, binding and crossing terms as it is explained in LoopSage (59). The term ℋ_rep_ represents the barrier effect imposed by replication forks, while ℋ_epi_ captures the spreading of epigenetic states, which may contribute to chromatin compartmentalization. In this paper, we adopt a Hamiltonian formalism, which enables us to explore the system’s preferred states by minimizing a generalized energy functional (104).

A key assumption of our model is that the distribution of states, 𝒮, follows a Boltzmann distribution:

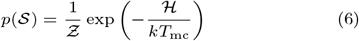

where *Ƶ* is the normalization constant, known in statistical physics as the *partition function*. The Boltzmann constant is set to *k* = 1 without loss of generality, while *T*_mc_ represents the *temperature* or *order parameter* of the stochastic system. Although *T*_mc_ plays a crucial role in system dynamics, its interpretation in biological systems remains elusive. In this study, we use *T*_mc_ to parameterize the system’s distribution: higher *T*_mc_ results in greater randomness, whereas lower *T*_mc_ leads to more ordered states. Assuming a Boltzmann distribution enables us to apply the well-established formalism of Monte Carlo methods (105).

Finally, we consider *N*_lef_ loop-extruding factors (LEFs), each defined by two degrees of freedom, (*m*_*i*_, *n*_*i*_), representing the link states of the graph, along with epigenetic states *s*_*i*_ for each monomer. As previously discussed, chromatin compartmentalization is a biophysical phenomenon associated with regions sharing similar epigenetic marks (17; 44; 45). In this study, we interpret compartmentalization as a *macrostate* in the sense of statistical physics, emerging from underlying *microstates* defined by these local epigenetic features (106). The system state is thus given by 𝒮 ≡ {*m*_*i*_, *n*_*i*_} ⊗ *s*_*k*_. This notation implicitly accounts for the time dependence of states, as typically encountered in Monte Carlo simulations: 𝒮(*t*) ≡ {*m*_*i*_(*t*), *n*_*i*_(*t*)} ⊗ *s*_*k*_(*t*). The graph 𝒢 consists of links either between consecutive nodes *i, i ±* 1 and between nodes *m*_*i*_ and *n*_*i*_ due to LEF-mediated interactions. Node states *s*_*k*_ can take five discrete values, {0, *±*1, *±*2}, which depend on the graph’s structure, determined by the distribution of links (Fig.S1(c)). The model’s resolution is set by the user-defined number of beads, *N*_beads_, ensuring that input data is downsampled to match this specified granularity.

### Loop Extrusion with LoopSage

Since RepliSage is a fork of LoopSage (59), we adopt the same energy function for modeling loop extrusion. However, to simplify the notation, we omit the explicit time dependence. Consequently, we can write down the following formula,

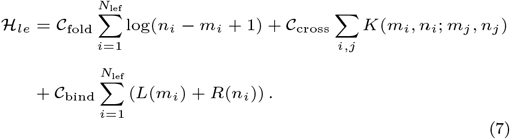

The meaning of this formula is that the energy is minimized in two ways: (i) by loop extrusion because of the first term (*C*_fold_ *<* 0), or (ii) by CTCF binding because of the last term (*C*_bind_ *<* 0). The potentials *L*(·) and *R*(·) are minimum in location where there is a loop in experimental data, and specifically a CTCF loop. The remaining term is the crossing energy, which is a penalty (*C*_cross_ *>* 0) in the case that two LEFs *i* and *j* cross each other *m*_*i*_ *< m*_*j*_ *< n*_*i*_ *< n*_*j*_. RepliSage allows for the testing of hypotheses involving multiple populations of loop extrusion factors (LEFs). In particular, it is possible to introduce a secondary population of LEFs, denoted *N*_lef,2_, characterized by a distinct folding factor *C*_fold,2_. This extension is especially relevant for modeling the mitotic phase of the cell cycle, which follows DNA replication. During mitosis, condensin complexes bind to chromatin and act as more processive LEFs, extruding longer loops at higher speeds compared to their interphase counterparts.

### Replication

The replication-related energy term, *H*_rep_, is a time-dependent component that models the interaction between loop extrusion factors (LEFs) and replication forks. Figure S1(f) illustrates several possible configurations of LEFs encountering replication forks. RepliSage assigns energetic penalties to configurations (iv) and (v), as these are physically forbidden due to the inability of LEFs to cross active replication forks. While configurations (ii) and (iii) are theoretically permissible, they are not expected to appear in equilibrium states of the system. This is because replication fork progression is significantly slower than loop extrusion, and the presence of such configurations would imply the existence of a mechanism by which LEFs routinely traverse replication forks - something that is not supported by current experimental evidence. Furthermore, we assume that the whole information of replication is embedded into the replication fraction function *f*_rep_(*x, t*) which is produced by a single run of a replication simulation and it is 1 if *x* is replicated at time *t* and 0 otherwise. Therefore, we can write,

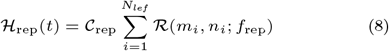

where the function ℛ(·) is a penalty that takes non-zero values in case that a cohesin crosses a replication fork or connects two distinct replicated locis,

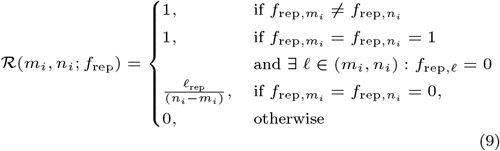

where *ℓ*_rep_ = |{*λ* ∈ (*m*_*i*_, *n*_*i*_) : *f*_*λ*_ *>* 1}| represents the number of replicated locis within the loop, and 𝒞_rep_ *>* 0 is a normalization term. The biological meaning of this term is that a replication fork acts as barrier for the motion of LEFs and thus if a LEF will be located close to a replication fork, it should prefer to stay there and not propagate further (Fig.S1(f). This term is particularly complicated because the replication fork trajectories (which are represented by *f*_rep_(*t*)) derived from Replikator are time-dependent as well, and therefore the energy landscape changes over time.

### Epigenetic Energy and Compartmentalization

Following the Potts model framework, each node in the graph represents a chromatin segment and can exist in one of five discrete epigenetic states, *s*_*i*_ ∈ {0, *±*1, *±*2}. These states correspond to different combinations of epigenetic marks (e.g., histone modifications or DNA accessibility), and can dynamically change over time. Biologically, regions of chromatin bearing similar epigenetic signatures tend to spatially associate, forming domains known as subcompartments (such as A1, A2, B1, B2 and B3 in Hi-C data). In our model, the alignment of multiple nodes in the same state increases their effective attraction, promoting the spatial segregation characteristic of compartmentalization. Therefore, we introduce five distinct states to capture a spectrum of possible chromatin identities and their contribution to 3D genome organization. Our model differs from the traditional Potts model, because it includes an additional epigenetic field which depends on the replication timing,

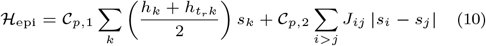

where *h*_*k*_ is an ‘*epigenetic field*’ (𝒞_*p*,1_ *<* 0) which is proportional to the replication timing of the current region, normalized to take values from −1 to 1 and assume that B compartment is linked to the negative values of the field. The term *h*_*t,k*_ represents the spread of the epigenetic state due to the propagation of the replication fork and it is,

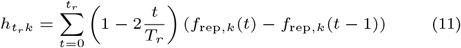

where *t*_*r*_ is the relative time since the replication process started, and *T*_*r*_ is the total duration of replication. Before the replication starts this epigenetic field is zero, and the average epigenetic field *h*_*k*_ is the dominant one. The term *h*_*t,k*_ captures the influence of replication forks on the epigenetic landscape during a specific stochastic simulation. It is positive when replication initiates (early replication) and negative when replication concludes (late replication). Biologically, *h*_*t,k*_ represents the potential of replication forks to reorganize epigenetic states. However, the precise mechanisms by which replication forks propagate or modify epigenetic marks remain incompletely understood, as highlighted in studies such as (39; 46; 47). To account for this uncertainty, RepliSage provides users with the option to disable this time-dependent effect by setting *h*_*t,k*_ ≡ *h*_*k*_, allowing expert users to tailor the model according to their domain-specific assumptions or hypotheses. The final term (*C*_*p*,2_ *>* 0) represents the interactions across the polymer graph assuming that 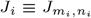 takes value 1 in locations of LEFs (*m*_*i*_(*t*), *n*_*i*_(*t*)), and zero in location where are no LEFs. Moreover, the interaction matrix is non-zero for sub-sequent beads *J*_*i,i*+1_ = 1. The difference |*s*_*i*_ − *s*_*j*_ | penalizes interactions of polymers with varying states, which leads to larger sizes of subcompartments.

### Parameter Balancing

When performing Monte Carlo simulations involving multiple energy components, it is important to understand how each term contributes to the model’s behavior. In RepliSage, quantifying the influence of individual energy terms is particularly challenging due to the lack of experimental data to constrain their relative weights. To address this, we propose an intuitive tuning strategy based on achieving a rough balance between energy contributions at equilibrium. In the following, we provide a set of default parameters that users can adopt with confidence, ensuring that the energy terms remain approximately comparable in magnitude under typical simulation conditions (Fig.S1(e)).

In the updated version of LoopSage, we slightly modified the model parametrization:

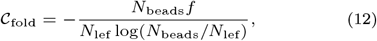

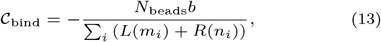

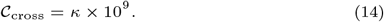

This re-parametrization ensures that when *f* = *b* = *κ* = 1, the energy terms are balanced, allowing for a more systematic tuning of parameters. The balancing is achieved by setting *C*_fold_ and *C*_bind_ to yield comparable equilibrium energies for *f* = *b* = 1, while *κ* takes high values so as to penalize improbable configurations of LEFs. The same principle is applied to subsequent energy terms.

Therefore, we can write for the replication and Potts model parameters,

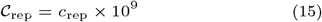

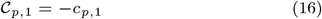

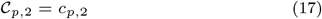

the choice of the sign for the Potts model is made in such a way that the energy is negative so as to be minimized when two interacting beads have the same node state. Note that we use positive values for *C*_*p*,2_ because we use the absolute value of the different of states |*s*_*i*_ − *s*_*j*_ | to define the interactions of Potts model. The replication term is multiplied by a large value of 10^9^ so as to penalize improbable configurations.

### The algorithm

To define the RepliSage algorithm, we decompose the energy change into two components: (i) terms associated with network rewiring, corresponding to LEF movements, and (ii) terms related to changes in epigenetic states. When a LEF move is proposed (i.e., modification of network links), two types of updates are possible: either a local random walk, shifting the LEF to neighboring positions (*m*_*i*_, *n*_*i*_) → (*m*_*i*_ *±* 1, *n*_*i*_ *±* 1), or a rebinding move, in which the LEF detaches and rebinds at a new random position with minimal loop length, i.e., *m*_*i*_ = *n*_*i*_. If an epigenetic update is proposed, the state *s*_*i*_ at a given node may transition to any state different from its current one. Therefore, we can write,

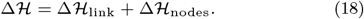

and thus we can easily handle the energies related with our Monte Carlo moves.

Therefore, the stochastic simulation of RepliSage can be summarized in the following steps,

1. The user defines the input <monospace>.bedpe </monospace>and replication curve data and from these data RepliSage determines the binding potentials and simulated replication fraction.
2. Initialize random initial configrations of LEFs (*m*_*i*_, *n*_*i*_) and states *s*_*i*_.
3. Initialize the matrix *J*_*ij*_ so as to be equal to 1 when *i* and *j* are connected by a LEF, and 0 otherwise.
4. We choose with probability *p*_1_ = 0.5 if it will be a node state change or rewiring move.
5. Compute the total energy difference Δℋ resulting from the proposed move, including any change in Potts interactions. Accept the move if Δℋ *<* 0, or with probability *e*^−Δ*H/kT*^ otherwise.
6. Change the interaction matrix if a rewiring move was accepted.
7. Repeat steps 4–6 for *N*_steps_ RepliSage epochs. In each epoch, perform *N*_sweep_ iterations, as specified by the user. To ensure adequate mixing of the system, it is generally recommended that *N*_sweep_ is comparable with the system size *N*_beads_, so that a substantial portion of the system’s states is proposed for change in each epoch.

In our Monte Carlo simulation we use sampling frequency *f*_mc_ so as to avoid correlated models, and a burnin period *τ*_burn-in_ so as to reach the equillibrium. The tricky part of RepliSage is that it is an out-of-equillibrium simulation in the sense that when the replication forks start their action, the system moves to a different state after the replication. Therefore, we expect that at least during replication RepliSage acts out of equillibrium.

### Molecular Dynamics

The molecular modeling approach assumes two polymer chains, each composed of *N*_beads_ monomers, where *N*_beads_ defines the granularity of the stochastic simulation. The total potential energy of the system is given by:

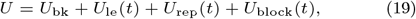

with each term representing a specific physical contribution. The backbone potential, *U*_bk_, includes strong covalent bonds between consecutive beads (*k*_bk_ = 5 × 10^5^ kJ*/*(mol · nm^2^)) with an equilibrium distance *d*_bk_ = 0.1 nm, angular forces (*k*_*ϕ*_ = 200 kJ*/*(mol · rad^2^), *θ* = *π*), and an excluded volume term acting only within the same chain:

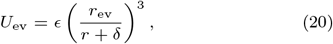

where *ϵ* = 200 kJ*/*(mol · nm^2^), *r*_ev_ = 0.1 nm, and *δ* = 0.01 nm. By setting *ϵ* = 0 at specific positions along the polymer chain, we model the activity of topoisomerases. This mechanism is crucial for the disentanglement and subsequent full separation of sister chromatids following the completion of the S phase.

The loop-formation potential, *U*_le_(*t*), is time-dependent and models LEF-mediated loop formation via harmonic bonds:

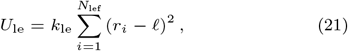

where *ℓ* = 0.1 nm is the equilibrium bond length and 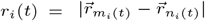 is the distance between LEF-bound beads. These interactions are weaker than backbone bonds, with strength *k*_le_ = 5 × 10 kJ*/*(mol · nm ^2^).

The replication potential, *U*_rep_(*t*), maintains strong harmonic bonds between paired monomers *i* and *i* + *N*_beads_ across the two chains prior to replication. As replication progresses, these bonds weaken, allowing chain separation:

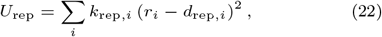

where both the bond strength and equilibrium distance between the two polymer chains depend on the replication state of each monomer *i*:

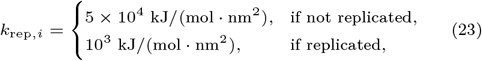

and,

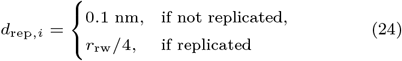

With this parameterization, DNA replication results in the gradual separation of the two polymer chains through the formation of long-range bonds, effectively modeling the de-attachment process. The factor 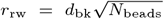 is used as an approximation of the characteristic length scale, based on the average distance of a random walk, which scales with the square root of the number of steps.

At the end of replication, upon entering the G2/M phase of the simulation, the replication-related potential *U*_rep_ is set to zero, and a new force is introduced to promote the spatial separation of sister chromatids. This force is modeled by the potential

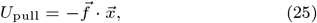

where 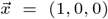 defines a fixed direction in space, and the magnitude of the force is *f* = *±*10^3^ kJ*/*(mol · nm^2^), with the sign depending on the polymer chain. This pulling force acts in competition with excluded volume interactions, facilitating chromatid disentanglement while preserving steric constraints.

The block-copolymer potential *U*_block_ is modeled as a Gaussian-shaped interaction (107–109), promoting attraction between monomers belonging to the same epigenetic compartment:

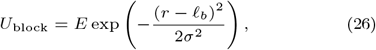

where the interaction strength *E* is defined as

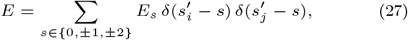

with *E*_*s*_ taking values specified in Table 1 when both interacting monomers belong to the same compartment, i.e., 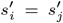. Here, *r* denotes the distance between monomers, *ℓ* _*b*_ is the preferred interaction distance, and *σ* sets the interaction range. The interaction energy *E*_*s*_ is stronger for the B compartment, which correlates with late replication timing. The range of excluded volume interactions is defined as *σ* = *r*_rw_*/*2, meaning that the interaction range is half the average distance of a random walk. Accordingly, the width of the Gaussian potential is set to 2*σ*. We assume that the equilibrium distance for compartment interactions is twice the loop equilibrium distance, *ℓ*_*b*_ = 0.2 nm, as compartmentalization reflects more distal, non-local interactions compared to loop extrusion. Compartment interactions are relatively weak, with A compartments considered more loose than B compartments (6; 110).

**Table 1.** Default OpenMM model parameters used in the molecular simulation, along with corresponding symbols, units, and values.

| Parameter Name | Symbol | Units | Value |
| --- | --- | --- | --- |
| Temperature | $T$ | Kelvin | 310 |
| Backbone bond strength | $k_{\text{bk}}$ | $\text{kJ}/(\text{mol} \cdot \text{nm}^2)$ | $5 \times 10^5$ |
| Backbone bond length | $d_{\text{bk}}$ | nm | 0.1 |
| Backbone angle force constant | $k_{\phi}$ | $\text{kJ}/(\text{mol} \cdot \text{rad}^2)$ | 200 |
| Backbone angle equilibrium | $\theta$ | rad | $\pi$ |
| Excluded volume strength | $\epsilon$ | $\text{kJ}/(\text{mol} \cdot \text{nm}^2)$ | 200 |
| Excluded volume radius | $r_{\text{ev}}$ | nm | 0.1 |
| Excluded volume shift | $\delta$ | nm | 0.01 |
| Excluded volume cut-off distance | $r_{\text{cut}}$ | nm | 0.2 |
| LEF bond strength | $k_{\text{le}}$ | $\text{kJ}/(\text{mol} \cdot \text{nm}^2)$ | $5 \times 10^4$ |
| LEF bond length | $\ell$ | nm | 0.1 |
| Replication bond strength | $k_{\text{rep}}$ | $\text{kJ}/(\text{mol} \cdot \text{nm}^2)$ | $5 \times 10^4$ |
| Replication bond strength (replicated) | $k_{\text{rep}}$ | $\text{kJ}/(\text{mol} \cdot \text{nm}^2)$ | $10^3$ |
| Replication bond length | $d_{\text{rep}}$ | nm | 0.1 |
| Replication bond length (replicated) | $d_{\text{rep}}$ | nm | $r_{\text{rw}}/4$ |
| Compartment interaction range | $\sigma$ | nm | $r_{\text{rw}}/2$ |
| Compartment equilibrium distance | $\ell_b$ | nm | 0.2 |
| Compartment energy (state $s = 2$) | $E_2$ | $\text{kJ}/(\text{mol} \cdot \text{nm}^2)$ | -0.2 |
| Compartment energy (state $s = 1$) | $E_1$ | $\text{kJ}/(\text{mol} \cdot \text{nm}^2)$ | -0.4 |
| Compartment energy (state $s = 0$) | $E_0$ | $\text{kJ}/(\text{mol} \cdot \text{nm}^2)$ | -0.6 |
| Compartment energy (state $s = -1$) | $E_{-1}$ | $\text{kJ}/(\text{mol} \cdot \text{nm}^2)$ | -0.8 |
| Compartment energy (state $s = -2$) | $E_{-2}$ | $\text{kJ}/(\text{mol} \cdot \text{nm}^2)$ | -1.0 |

### Simulation of Cell Cycle

To model the cell cycle, we divided the simulation into three distinct phases (G1, S, and G2/M) each requiring specific energy modifications and implementation strategies. In the G1 phase, chromatin dynamics are driven by loop extrusion, where CTCF proteins act as static barriers to loop extrusion factors (LEFs), and epigenetic states evolve via a Potts model. The number of LEFs in this phase was set to *N*_lef_ = 1000, with a baseline extrusion speed *f*_1_ = 1. In the S phase, replication forks, imported from the *Replikator* module, are introduced and act as dynamic, transient obstacles to LEF progression. Once replication is complete, the two daughter DNA chains remain closely associated. The G2/M phase is modeled by (i) introducing a second population of LEFs (*N*_lef,2_ = 1000) with a higher extrusion rate *f*_2_ = 2, representing condensins, and (ii) enabling local relaxation of excluded volume interactions through random Monte Carlo moves, mimicking topoisomerase activity with probability *p*_top_ = 0.01. This reflects experimental findings that condensins operate 2–5 times faster than cohesins and can bypass them without displacement (111), and that topoisomerases act throughout the cell cycle. Both LEF populations operate independently and continuously extrude loops. Chromosome-wide simulations were performed using a coarse-grained polymer model of two chains, each consisting of 10,000 beads. The Monte Carlo order parameter was fixed at *T*_mc_ = 1.8 to balance structural order and fluctuation. Simulations ran for 2 × 10^5^ steps using the Metropolis algorithm, with configurations sampled every *f*_mc_ = 200 steps after a burn-in period of 1,000 steps. The simulation phases were segmented as follows: G1 (0– 5 × 10^4^ steps), S phase (5 × 10^4^–10^5^), and G2/M (10^5^–2 × 10^5^ steps). Execution of both the stochastic Monte Carlo simulation and associated molecular dynamics via OpenMM required slightly less than one day on standard computational infrastructure.

### Parameter Study Setup

To explore how model parameters influence the relationship between DNA replication and chromatin structure, we performed a targeted parameter study on a representative region of human chromosome 14, spanning genomic coordinates [70835000, 98674700]. This specific segment was selected to balance biological relevance and computational efficiency, as simulating an entire chromosome at high resolution would be computationally prohibitive.

Simulations were carried out using two polymer chains, each comprising *N*_beads_ = 2000 beads. The total simulation time was set to *N*_steps_ = 2 × 10^5^ Monte Carlo steps. The system was first equilibrated during the G1 phase for *τ*_rep_ = 5 × 10^4^ steps. Replication was then simulated over *T*_*r*_ = 5 × 10^4^ steps (S phase), followed by additional steps to allow the system to reach equilibrium post-replication (G2/M phase). We mainly focused on the behavior of the system under the change of the order parameter taking the following values *T*_mc_ ∈ [1, 1.4, 1.8, 2.2, 2.6, 3.0]. This controlled setup enabled systematic investigation of parameter-dependent chromatin reorganization across the cell cycle. For each temperature, RepliSage was independently executed 10 times, and the statistical averages from these runs were presented in Fig.6.

### Definition of the Structural Metrics

In this study, we analyzed several structural metrics that characterize chromatin folding. For completeness and clarity, we provide formal definitions of these metrics below.

- **Radius of gyration (***R*_*g*_): Quantifies the spatial extent of the chromatin polymer by measuring the mean squared distance of monomers from the center of mass:

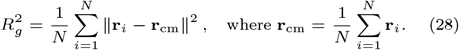
- **Convex hull volume (***V*_**convex**_): Represents the 3D volume enclosing all monomer positions, calculated as:

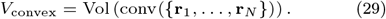

Computed using the scipy.spatial.ConvexHull class, which decomposes the hull into tetrahedra and sums their volumes:

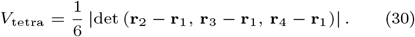
- **Ellipsoid volume:** The ellipsoid volume estimates the effective 3D space occupied by the chromatin conformation, calculated using the eigenvalues *λ*_1_, *λ*_2_, *λ*_3_ of the covariance matrix of the coordinates:

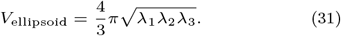
- **Ellipsoid axis ratio:** This ratio characterizes elongation of the structure. It is defined as the ratio between the largest and smallest principal axes of the fitted ellipsoid:

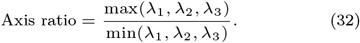
- **Asphericity:** Asphericity quantifies the deviation from spherical symmetry. A perfectly spherical object has *A* = 0:

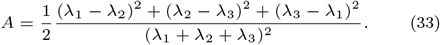
- **Anisotropy:** Anisotropy measures directional preference in spatial structure, with values closer to 1 indicating stronger bias:

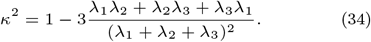
- **Mean distance to the center of mass:** This metric reflects how spread out the structure is around its center:

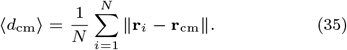
- **Maximum distance to the center of mass:** Indicates the furthest extent of the structure from its geometric center:

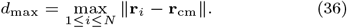
- **Mean nearest-neighbor distance:** Describes local density by averaging the distance to the closest neighboring bead for each monomer:

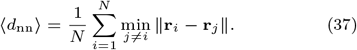
- **Bounding box volume:** The volume of the axis-aligned bounding box enclosing the structure, defined by extremal coordinates:

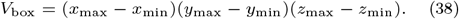
- **End-to-end distance:** Measures the linear span of the polymer from its first to last bead:

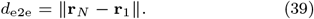
- **Contour length:** The total length along the polymer path, summing distances between consecutive beads:

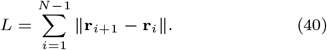
- **Straightness index:** Captures the degree of linearity of the polymer, computed as the ratio of end-to-end distance to contour length:

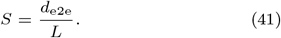

A value close to 1 indicates a straight configuration, whereas lower values suggest coiling or looping.

### Parallelization

Monte Carlo simulations, particularly those based on the Metropolis algorithm, are computationally demanding due to the large number of stochastic updates required to explore the energy landscape thoroughly. Nevertheless, their key advantage lies in generating structural ensembles rather than relying on single averaged configurations, allowing more accurate representation of biological variability.

To improve computational efficiency, RepliSage incorporates just-in-time (JIT) compilation using the `numba` Python library (112), which translates critical parts of the Monte Carlo code into low-level machine instructions. This optimization results in substantial speedups, with performance gains of up to 100-fold as reported in the library’s documentation. The same approach has also been applied to develop an optimized version of LoopSage. For the molecular dynamics component, RepliSage employs OpenMM (102; 103), which supports parallel execution on multi-core CPUs and enables GPU acceleration via OpenCL or CUDA backends. The complete RepliSage pipeline has been validated across a range of computational environments, including personal laptops, desktop workstations, and high-performance computing (HPC) clusters.

## Supplementary Figures

**Fig. S1.**
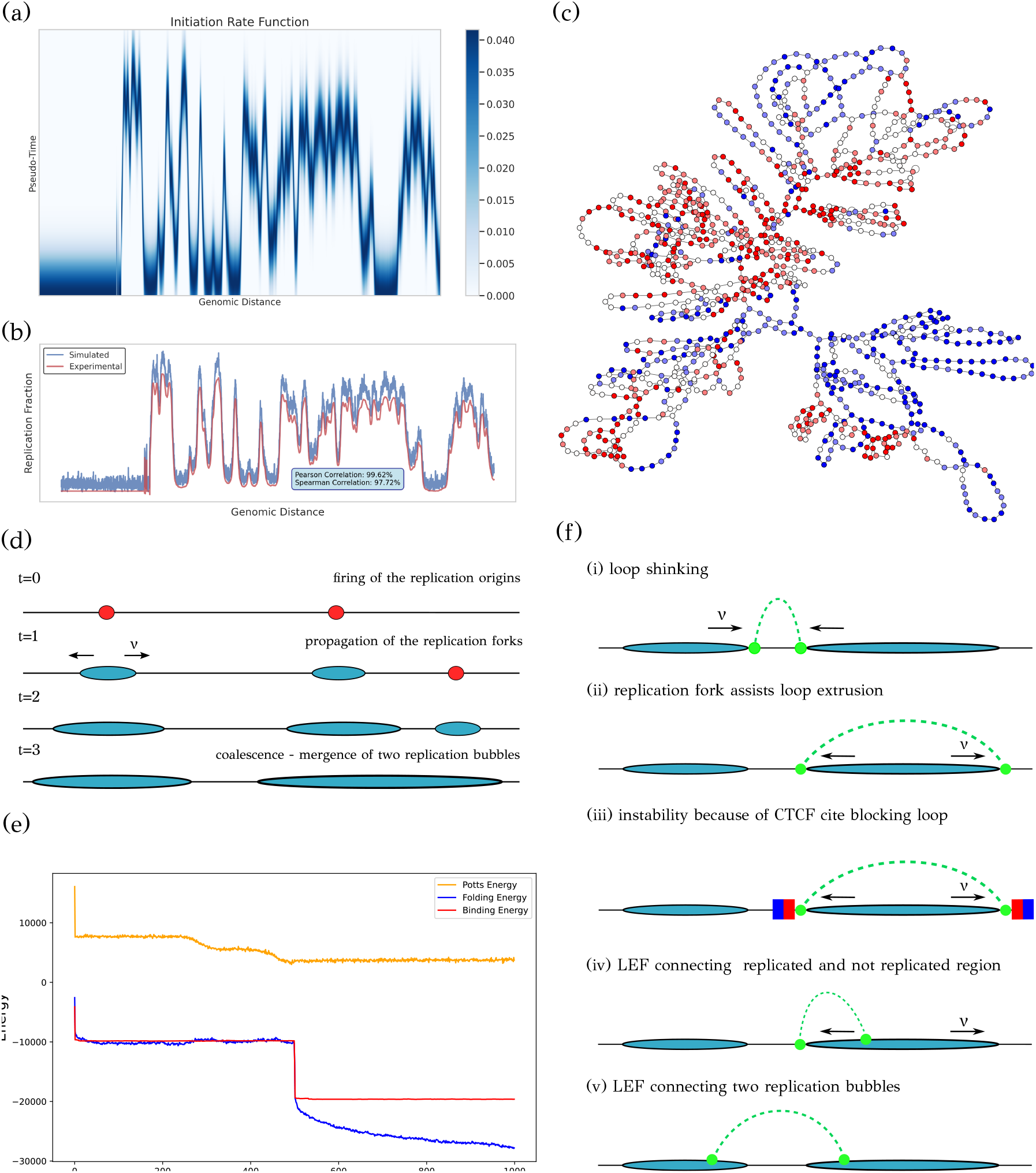
Overview of the RepliSage methodology. (a) The initiation rate function *I*(*x, t*) follows a Gaussian distribution centered on the replication timing *µ*(*x*), defining the probability of origin firing over time. (b) Replication simulations produce average replication fraction curves highly correlated with experimental data (Pearson 99.62%, Spearman 97.72%, based on 100 trials), validating the model’s ability to capture fork dynamics. (c) Example chromatin graph where node colors indicate epigenetic states (blue: negative, red: positive), and edges represent either sequential or loop-forming LEF-mediated connections. These graphs define microstates used in stochastic sampling, and can be translated to 3D structures using OpenMM with harmonic bonds (edges) and block-copolymer interactions (nodes). (d) Schematic of stochastic replication: origins fire with probability *I*(*x, t*), generating bidirectional forks; replication completes upon full coverage. (e) Interplay of RepliSage energy components: loop extrusion and binding terms interact, while the Potts epigenetic term is minimized independently due to opposing sign. (f) Schematic of LEF-fork compatibility: replication forks act as moving barriers. LEFs are only allowed between regions that are either fully replicated or unreplicated; all configurations that require LEFs to span active forks are disallowed.

**Fig. S2.**
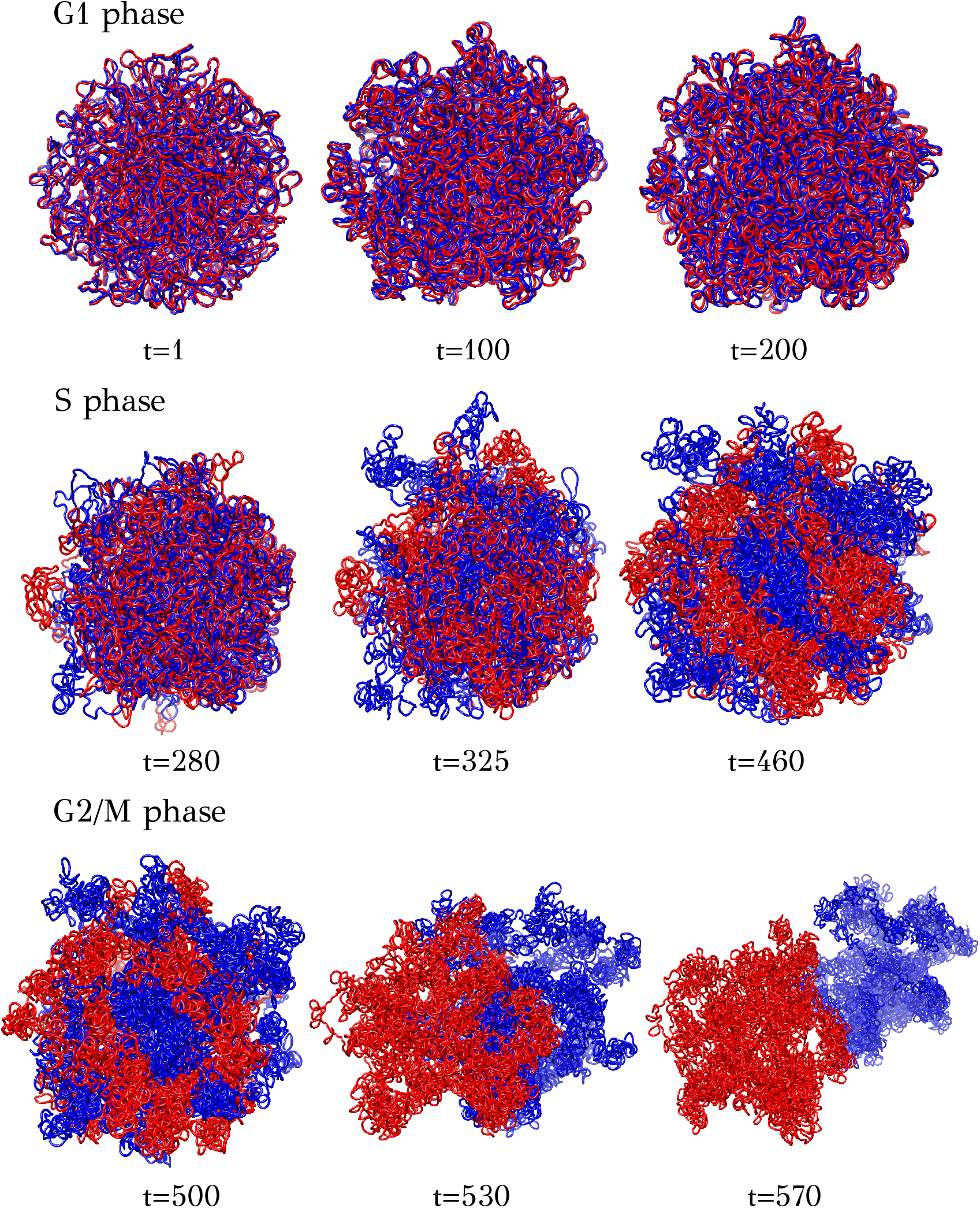
Selected frames from a RepliSage simulation of regular replication for the GM12878 dataset. In the G1 phase, the chromatin adopts a compact, globular conformation with the two polymer chains attached. During the S phase, replication forks initiate and propagate, leading to gradual de-attachment of the chains. By the end of S phase, the chains are mostly separated but not yet fully distinct. In the G2/M phase, longer loops form, giving rise to a supercoiled structure and promoting full separation of the chains. Final compaction follows upon complete separation of the replicated chromosomes.

**Fig. S3.**
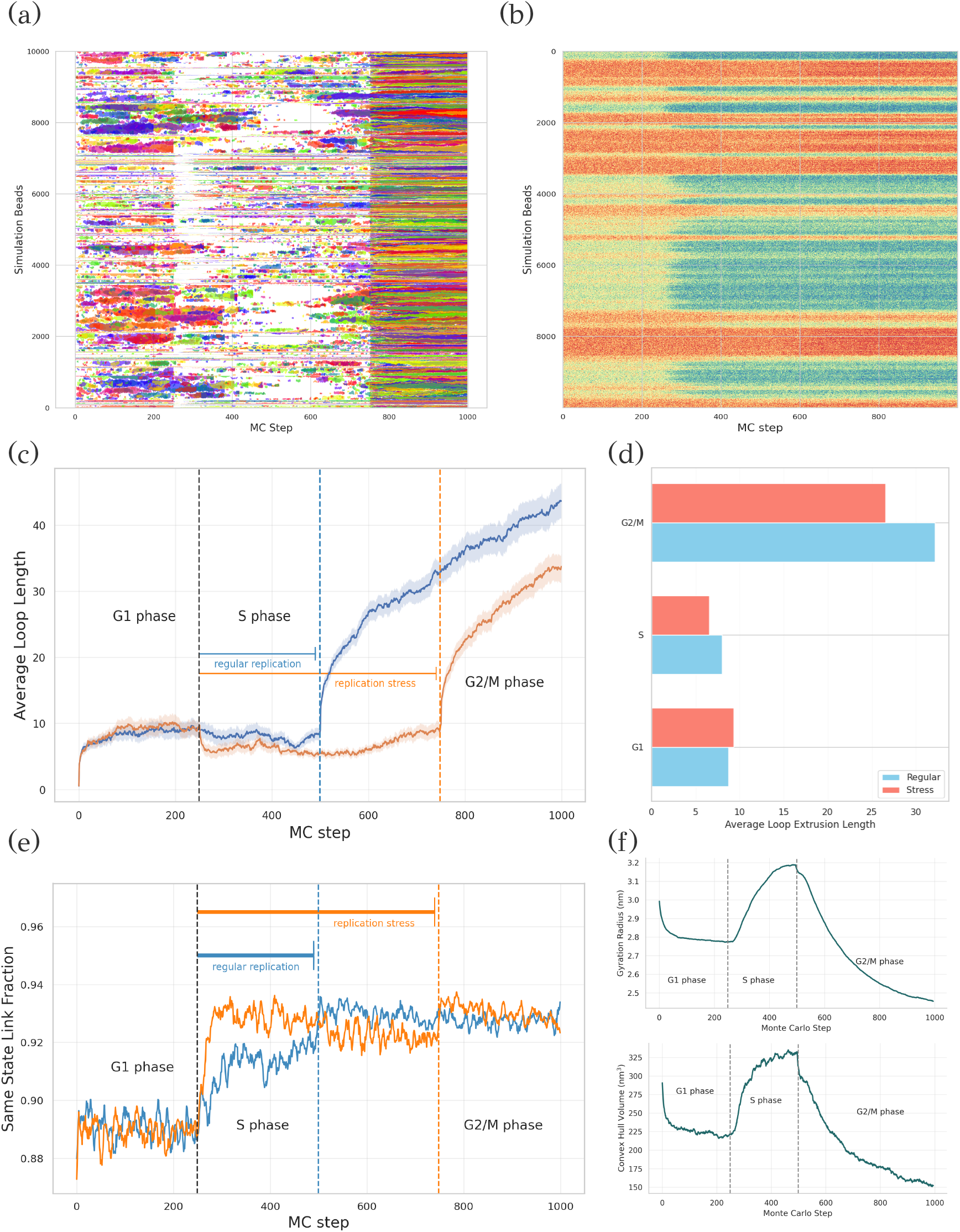
Simulation results for chromosome 14 using CTCF ChIA-PET data from the GM12878 cell line. (a) LEF trajectories, with each extruded loop shown in a different color. During the S phase, many loops disassemble into smaller ones due to replication forks acting as moving barriers. In the G2/M phase, condensins drive the formation of longer loops. (b) Distribution and reorganization of epigenetic states during the S phase. (c) Time-resolved average loop length across cell cycle phases, comparing regular and stress-induced replication. (d) Phase-specific average loop lengths. (e) Fraction of links connecting nodes with identical epigenetic states under regular and stress conditions. (f) Chromatin compaction during the cell cycle, quantified using the radius of gyration and convex hull volume.

**Fig. S4.**
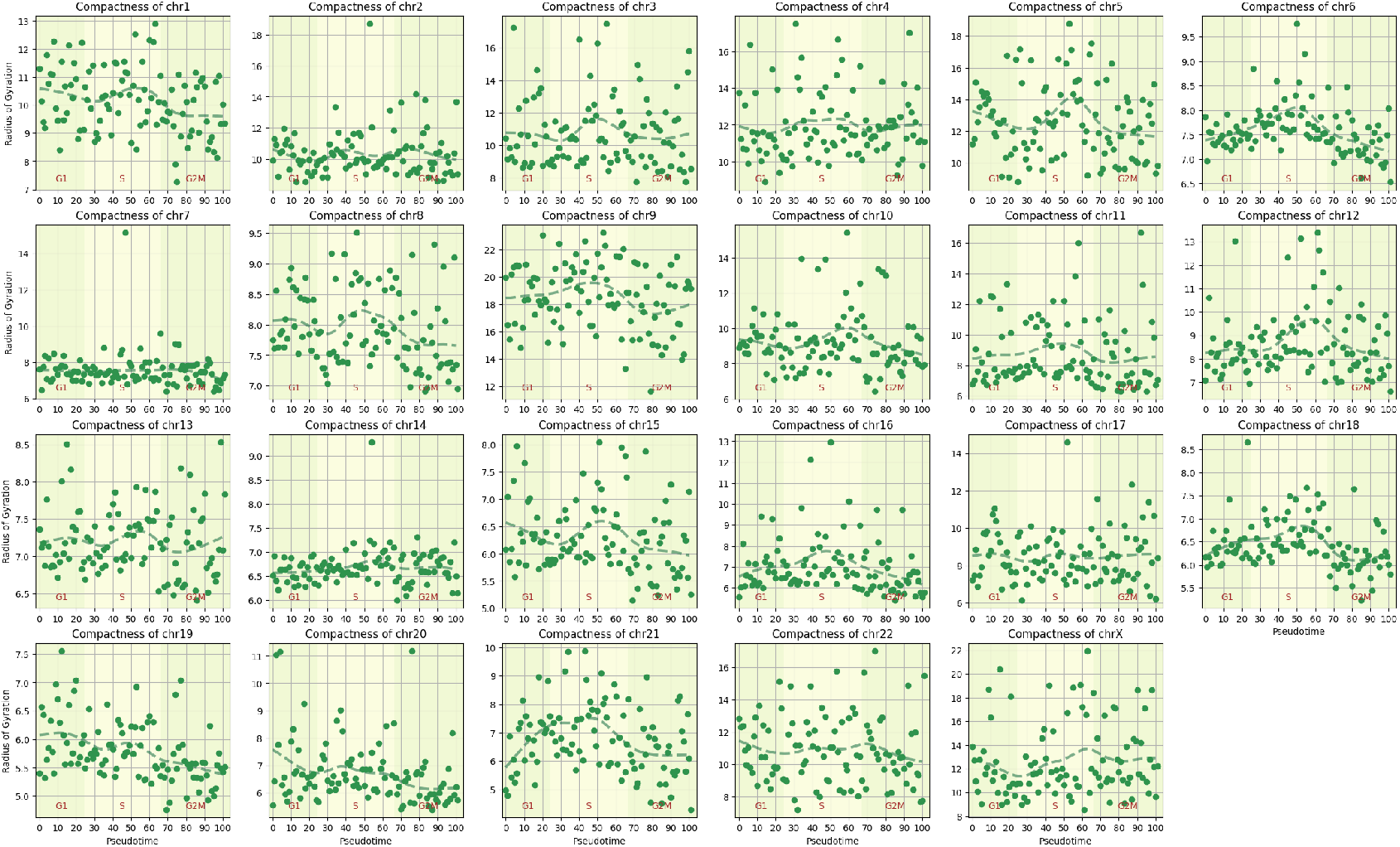
Radius of Gyration Measurements from NucDynamics Simulations on K562 Data. Radius of gyration values were obtained from NucDynamics simulations based on the K562 dataset from (82). Each point represents a specific metacell at a given pseudotime along the cell cycle trajectory, corresponding to a single chromosome. LOESS (Locally Estimated Scatterplot Smoothing) curves are overlaid to illustrate general trends in chromosomal compaction over time.

**Fig. S5.**
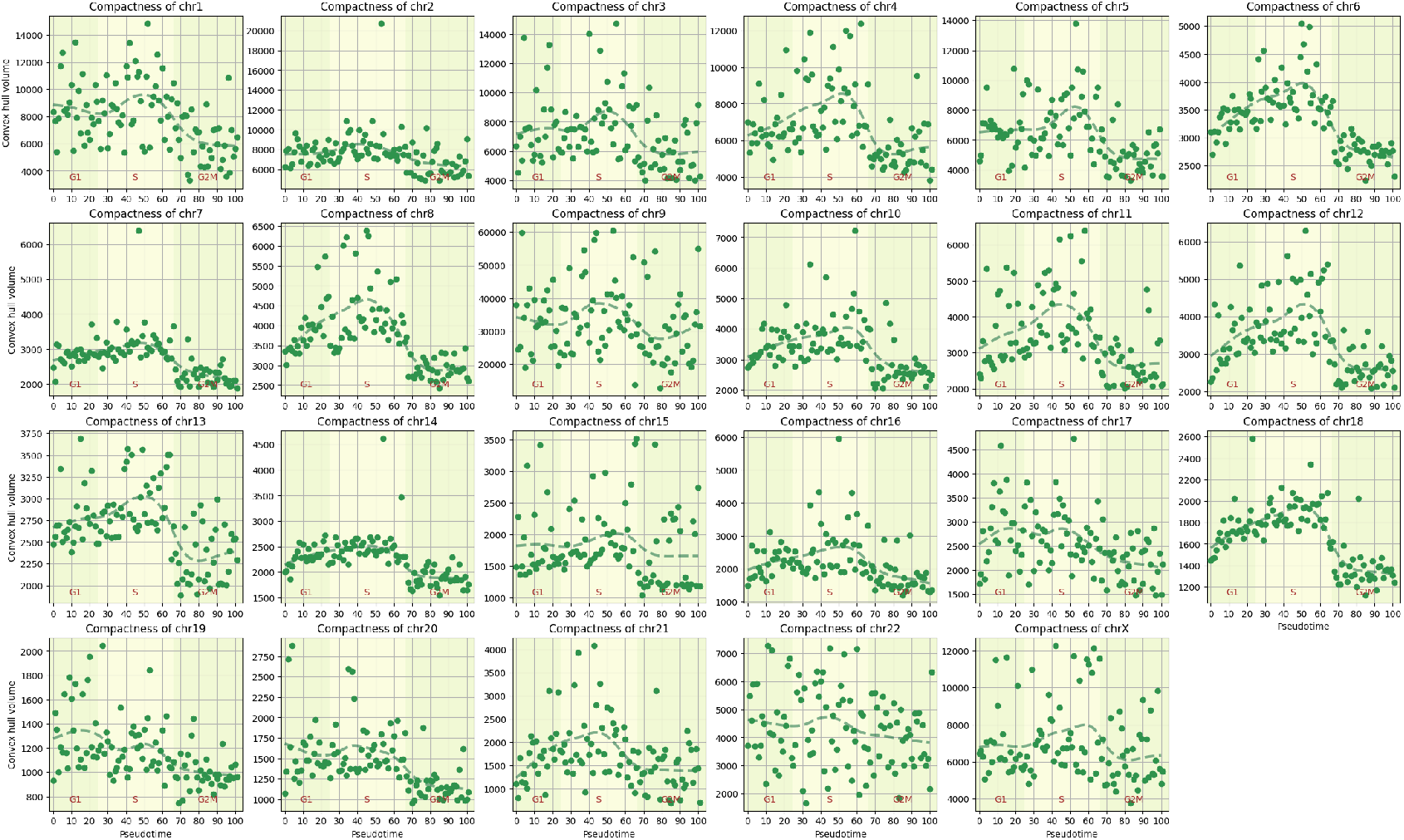
Convex hull volume measurements from NucDynamics Simulations on K562 Data. Convex hull volume values were obtained from NucDynamics simulations based on the K562 dataset from (82). Each point represents a specific metacell at a given pseudotime along the cell cycle trajectory, corresponding to a single chromosome. LOESS (Locally Estimated Scatterplot Smoothing) curves are overlaid to illustrate general trends in chromosomal compaction over time.

**Fig. S6.**
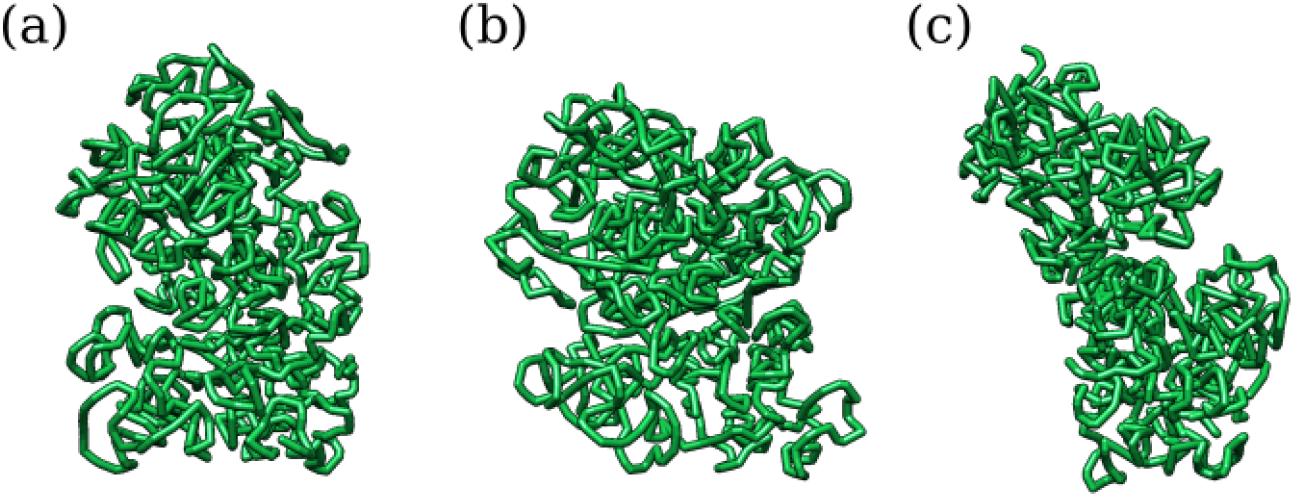
Representative NucDynamics structures of the human chromosome 14 modelled on K562 metacells from (82). (a) Representative structure of chr14 from the G1 phase (metacell 10). (b) Representative structure of chr14 from the S phase (metacell 42). (c) Representative structure of chr14 from the G2M phase (metacell 91).

**Fig. S7.**
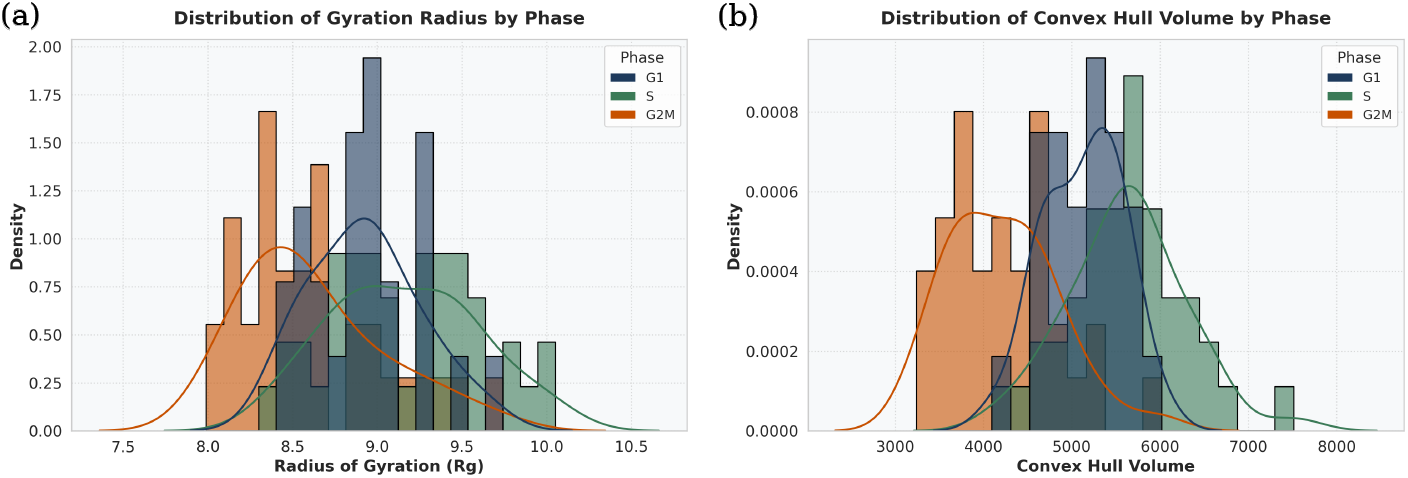
Comparison between phases for the NucDynamics structures of the K562 cell line. (a) Radius of Gyration measurements. (b) Convex Hull volume measurements. Each data point represents the mean value across all chromosomes at a specific pseudotime during cell cycle progression.

**Fig. S8.**
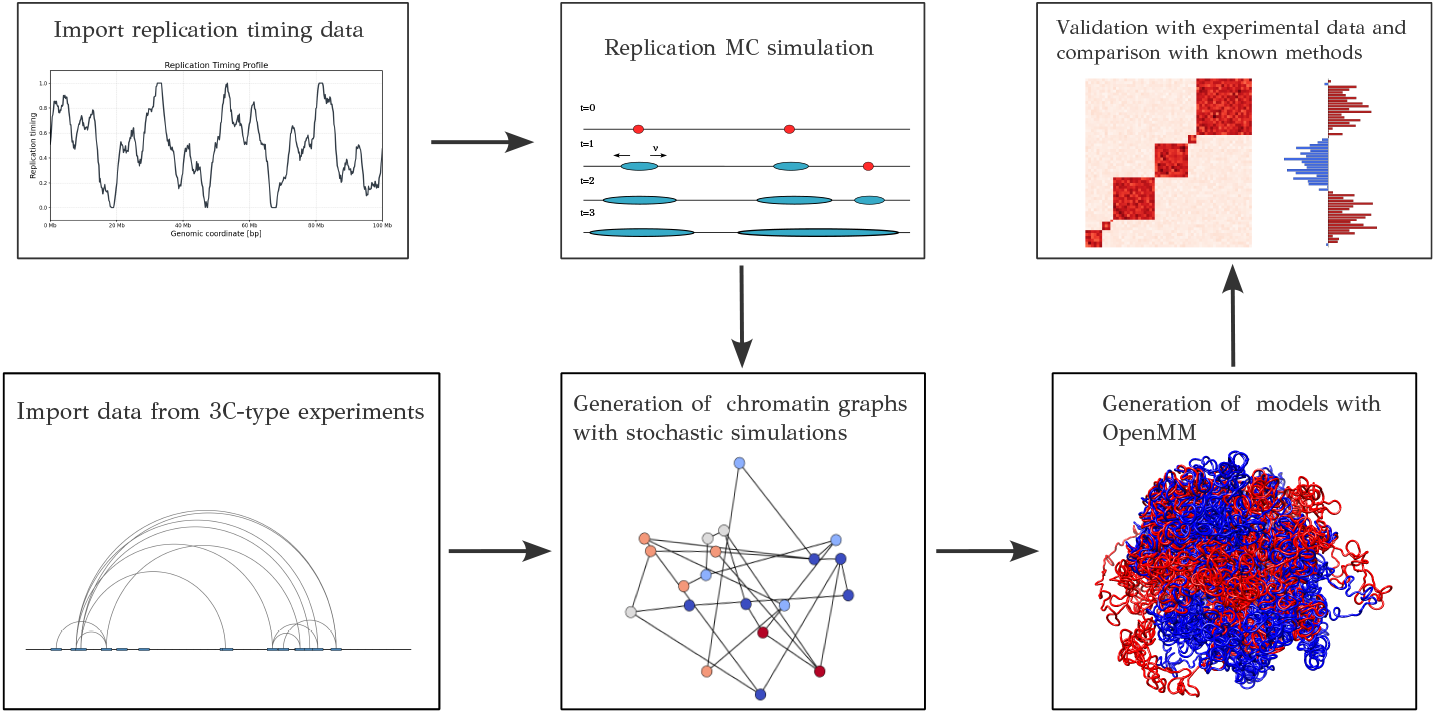
General scheme of the RepliSage model (*graphical abstract*). Replication timing data are imported, pre-processed, and passed to the Monte Carlo replication simulation module (*Replikator*) to simulate origin activation and replication fork propagation. In parallel, RepliSage imports a user-provided bedpe file from a 3C-type experiment describing chromatin loops (in this study, CTCF ChIA-PET data were used). The trajectories of replication forks and chromatin loops are then passed to the stochastic energy optimization module of RepliSage, which performs Metropolis sampling to generate graph states where links represent backbone bonds and loop extrusion factor (LEF) mediated links, and node states represent epigenetic states. The stochastic simulation applies non-equilibrium rules to simulate all phases of the cell cycle. The generated time-dependent states can be further imported into OpenMM for force-field optimization to reconstruct the 3D structural trajectory. Finally, the resulting structures are compared to established models, including single-cell 3D reconstructions, and validated against orthogonal experimental datasets.

**Fig. S9.**
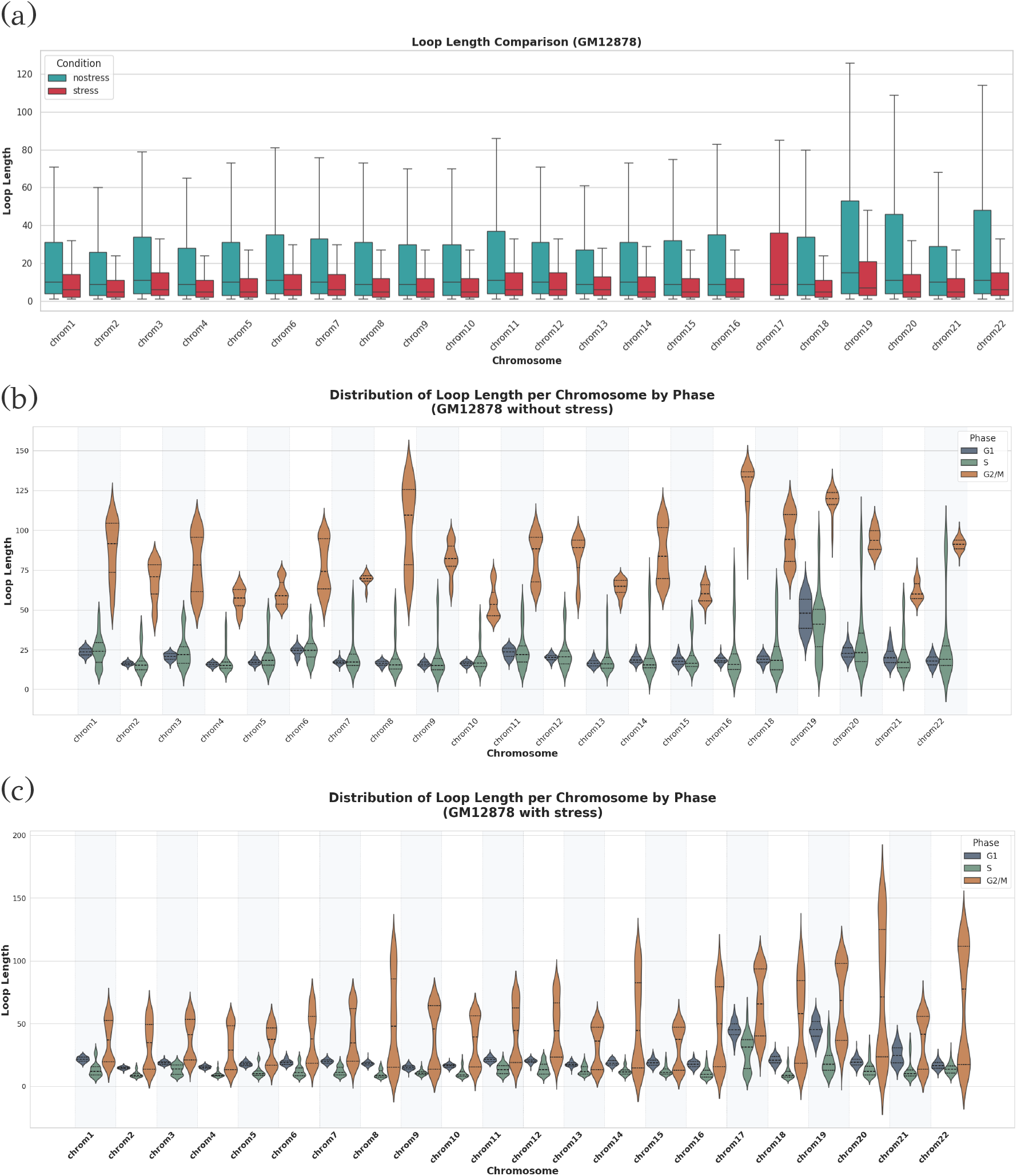
Comparison of loop length distributions across cell cycle phases using RepliSage simulations based on GM12878 CTCF ChIA-PET data, applied to 100 kb-resolution models of all chromosomes. (a) Loop length distributions per chromosome under regular replication versus replication stress conditions. (b) Phase-specific loop length distributions for each chromosome under regular replication. (c) Phase-specific loop length distributions for each chromosome under replication stress conditions.

**Fig. S10.**
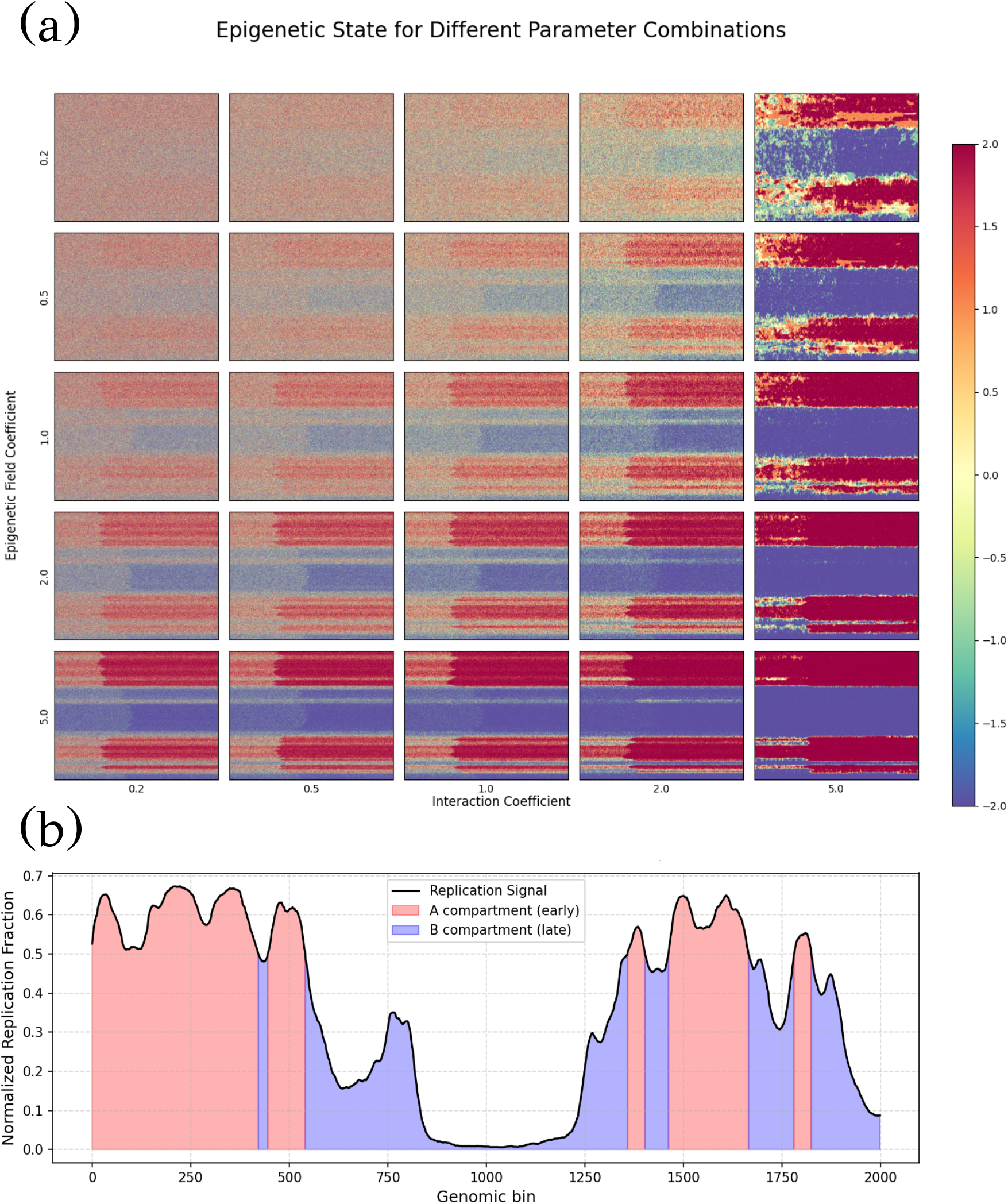
Balance of Potts energy terms in the RepliSage Hamiltonian. (a) Trajectories of epigenetic states over Monte Carlo time for different combinations of field and interaction coefficients, *C*_*p*,1_ and *C*_*p*,2_. The x-axis represents Monte Carlo time, and the y-axis shows epigenetic states across network nodes, ordered sequentially. (b) Replication timing data for the modeled region [70835000, 98674700] of chromosome 14 (GM12878, CTCF ChIA-PET). Early-replicating domains correspond to positive epigenetic states, typically associated with A compartments. As shown in (a), such compartment-like collective behavior emerges when both field and interaction terms are sufficiently strong.

## Footnotes

1 https://satijalab.org/seurat/articles/cell_cycle_vignette.html

## References

1. A. S. Hansen, “Ctcf as a boundary factor for cohesin-mediated loop extrusion: evidence for a multi-step mechanism,” Nucleus, vol. 11, no. 1, pp. 132–148, 2020. PMID: 32631111.

2. A. Agarwal, S. Korsak, A. Choudhury, and D. Plewczynski, “The dynamic role of cohesin in maintaining human genome architecture,” BioEssays, vol. 45, no. 10, p. 2200240, 2023.

3. M. Lazniewski, W. K. Dawson, A. M. Rusek, and D. Plewczynski, “One protein to rule them all: The role of ccctc-binding factor in shaping human genome in health and disease,” Seminars in Cell and Developmental Biology, vol. 90, pp. 114–127, 2019. 3D Genome and Diseases.

4. K. Rippe, “Liquid–liquid phase separation in chromatin,” Cold Spring Harbor perspectives in biology, vol. 14, no. 2, p. a040683, 2022.

5. F. Erdel and K. Rippe, “Formation of chromatin subcompartments by phase separation,” Biophysical journal, vol. 114, no. 10, pp. 2262–2270, 2018.

6. J.-P. Fortin and K. D. Hansen, “Reconstructing a/b compartments as revealed by hi-c using long-range correlations in epigenetic data,” Genome biology, vol. 16, pp. 1–23, 2015.

7. H. L. Harris, H. Gu, M. Olshansky, A. Wang, I. Farabella, Y. Eliaz, A. Kalluchi, A. Krishna, M. Jacobs, G. Cauer, et al., “Chromatin alternates between a and b compartments at kilobase scale for subgenic organization,” Nature communications, vol. 14, no. 1, p. 3303, 2023.

8. T. Cremer and M. Cremer, “Chromosome territories,” Cold Spring Harbor perspectives in biology, vol. 2, no. 3, p. a003889, 2010.

9. G. Ozer, A. Luque, and T. Schlick, “The chromatin fiber: multiscale problems and approaches,” Current opinion in structural biology, vol. 31, pp. 124–139, 2015.

10. C. Weinreb and B. J. Raphael, “Identification of hierarchical chromatin domains,” Bioinformatics, vol. 32, no. 11, pp. 1601–1609, 2016.

11. G. Li and D. Reinberg, “Chromatin higher-order structures and gene regulation,” Current opinion in genetics & development, vol. 21, no. 2, pp. 175–186, 2011.

12. C. L. Woodcock and R. P. Ghosh, “Chromatin higher-order structure and dynamics,” Cold Spring Harbor perspectives in biology, vol. 2, no. 5, p. a000596, 2010.

13. S. Korsak, K. Banecki, and D. Plewczynski, “Multiscale molecular modeling of chromatin with multimm: From nucleosomes to the whole genome,” Computational and Structural Biotechnology Journal, vol. 23, pp. 3537–3548, 2024.

14. B. Fierz and M. G. Poirier, “Biophysics of chromatin dynamics,” Annual review of biophysics, vol. 48, no. 1, pp. 321–345, 2019.

15. M. R. Hübner and D. L. Spector, “Chromatin dynamics,” Annual review of biophysics, vol. 39, no. 1, pp. 471–489, 2010.

16. E. I. Prieto and K. Maeshima, “Dynamic chromatin organization in the cell,” Essays in biochemistry, vol. 63, no. 1, pp. 133–145, 2019.

17. M. Di Pierro, R. R. Cheng, E. Lieberman Aiden, P. G. Wolynes, and J. N. Onuchic, “De novo prediction of human chromosome structures: Epigenetic marking patterns encode genome architecture,” Proceedings of the National Academy of Sciences, vol. 114, no. 46, pp. 12126–12131, 2017.

18. H. Zhang, X.-J. Tian, A. Mukhopadhyay, K. Kim, and J. Xing, “Statistical mechanics model for the dynamics of collective epigenetic histone modification,” Physical review letters, vol. 112, no. 6, p. 068101, 2014.

19. Q. Mo and F. Liang, “A hidden ising model for chip-chip data analysis,” Bioinformatics, vol. 26, no. 6, pp. 777–783, 2010.

20. Q. Mo and F. Liang, “Bayesian modeling of chip-chip data through a high-order ising model,” Biometrics, vol. 66, no. 4, pp. 1284–1294, 2010.

21. D. Michieletto, E. Orlandini, and D. Marenduzzo, “Polymer model with epigenetic recoloring reveals a pathway for the de novo establishment and 3d organization of chromatin domains,” Physical Review X, vol. 6, no. 4, p. 041047, 2016.

22. K. Adachi and K. Kawaguchi, “Chromatin state switching in a polymer model with mark-conformation coupling,” Physical Review E, vol. 100, no. 6, p. 060401, 2019.

23. B. M. McCoy and T. T. Wu, The Two-Dimensional Ising Model. Cambridge, MA and London, England: Harvard University Press, 1973.

24. L. Beaudin, “A review of the potts model,” Rose-Hulman Undergraduate Mathematics Journal, vol. 8, no. 1, p. 13, 2007.

25. Y. Hu and B. Stillman, “Origins of dna replication in eukaryotes,” Molecular cell, vol. 83, no. 3, pp. 352–372, 2023.

26. A. Costa and J. F. Diffley, “The initiation of eukaryotic dna replication,” Annual review of biochemistry, vol. 91, no. 1, pp. 107–131, 2022.

27. D. McIntosh and J. J. Blow, “Dormant origins, the licensing checkpoint, and the response to replicative stresses,” Cold Spring Harbor perspectives in biology, vol. 4, no. 10, p. a012955, 2012.

28. M. Méchali, “Eukaryotic dna replication origins: many choices for appropriate answers,” Nature reviews Molecular cell biology, vol. 11, no. 10, pp. 728–738, 2010.

29. S. Kagiwada, K. Kurimoto, T. Hirota, M. Yamaji, and M. Saitou, “Replication-coupled passive dna demethylation for the erasure of genome imprints in mice,” The EMBO journal, vol. 32, no. 3, pp. 340–353, 2013.

30. C. Conti, B. Saccà, J. Herrick, C. Lalou, Y. Pommier, and A. Bensimon, “Replication fork velocities at adjacent replication origins are coordinately modified during dna replication in human cells,” Molecular biology of the cell, vol. 18, no. 8, pp. 3059–3067, 2007.

31. R. Milo and R. Phillips, Cell biology by the numbers. Garland Science, 2015.

32. F. Corsi, E. Rusch, and A. Goloborodko, “Loop extrusion rules: the next generation,” Current Opinion in Genetics & Development, vol. 81, p. 102061, 2023.

33. M. Ganji, I. A. Shaltiel, S. Bisht, E. Kim, A. Kalichava, C. H. Haering, and C. Dekker, “Real-time imaging of dna loop extrusion by condensin,” Science, vol. 360, no. 6384, pp. 102–105, 2018.

34. T. Nagano, Y. Lubling, C. Várnai, C. Dudley, W. Leung, Y. Baran, N. Mendelson Cohen, S. Wingett, P. Fraser, and A. Tanay, “Cell-cycle dynamics of chromosomal organization at single-cell resolution,” Nature, vol. 547, no. 7661, pp. 61–67, 2017.

35. M. Minamino, C. Bouchoux, B. Canal, J. F. Diffley, and F. Uhlmann, “A replication fork determinant for the establishment of sister chromatid cohesion,” Cell, vol. 186, no. 4, pp. 837–849, 2023.

36. M.-E. Terret, R. Sherwood, S. Rahman, J. Qin, and P. V. Jallepalli, “Cohesin acetylation speeds the replication fork,” Nature, vol. 462, no. 7270, pp. 231–234, 2009.

37. S. Villa-Hernández and R. Bermejo, “Cohesin dynamic association to chromatin and interfacing with replication forks in genome integrity maintenance,” Current genetics, vol. 64, no. 5, pp. 1005–1013, 2018.

38. H. Bierne and B. Michel, “When replication forks stop,” Molecular microbiology, vol. 13, no. 1, pp. 17–23, 1994.

39. C. Alabert and A. Groth, “Chromatin replication and epigenome maintenance,” Nature reviews Molecular cell biology, vol. 13, no. 3, pp. 153–167, 2012.

40. C. Marchal, J. Sima, and D. M. Gilbert, “Control of dna replication timing in the 3d genome,” Nature Reviews Molecular Cell Biology, vol. 20, no. 12, pp. 721–737, 2019.

41. P. A. Zhao, J. C. Rivera-Mulia, and D. M. Gilbert, “Replication domains: genome compartmentalization into functional replication units,” DNA replication: from old principles to new discoveries, pp. 229–257, 2017.

42. T. Ryba, I. Hiratani, J. Lu, M. Itoh, M. Kulik, J. Zhang, T. C. Schulz, A. J. Robins, S. Dalton, and D. M. Gilbert, “Evolutionarily conserved replication timing profiles predict long-range chromatin interactions and distinguish closely related cell types,” Genome research, vol. 20, no. 6, pp. 761–770, 2010.

43. B. Moindrot, B. Audit, P. Klous, A. Baker, C. Thermes, W. de Laat, P. Bouvet, F. Mongelard, and A. Arneodo, “3d chromatin conformation correlates with replication timing and is conserved in resting cells,” Nucleic acids research, vol. 40, no. 19, pp. 9470–9481, 2012.

44. E. M. Hildebrand and J. Dekker, “Mechanisms and functions of chromosome compartmentalization,” Trends in biochemical sciences, vol. 45, no. 5, pp. 385–396, 2020.

45. P. B. Singh and A. G. Newman, “On the relations of phase separation and hi-c maps to epigenetics,” Royal Society open science, vol. 7, no. 3, p. 191976, 2020.

46. S. J. Charlton, V. Flury, Y. Kanoh, A. V. Genzor, L. Kollenstart, W. Ao, P. Brøgger, M. B. Weisser, M. Adamus, N. Alcaraz, et al., “The fork protection complex promotes parental histone recycling and epigenetic memory,” Cell, vol. 187, no. 18, pp. 5029–5047, 2024.

47. I. Whitehouse and D. J. Smith, “Chromatin dynamics at the replication fork: there’s more to life than histones,” Current opinion in genetics & development, vol. 23, no. 2, pp. 140–146, 2013.

48. J. H. Gibcus, K. Samejima, A. Goloborodko, I. Samejima, N. Naumova, J. Nuebler, M. T. Kanemaki, L. Xie, J. R. Paulson, W. C. Earnshaw, et al., “A pathway for mitotic chromosome formation,” Science, vol. 359, no. 6376, p. eaao6135, 2018.

49. K. Nasmyth, “Disseminating the genome: joining, resolving, and separating sister chromatids during mitosis and meiosis,” Annual review of genetics, vol. 35, no. 1, pp. 673–745, 2001.

50. J. L. Nitiss, “Dna topoisomerase ii and its growing repertoire of biological functions,” Nature Reviews Cancer, vol. 9, no. 5, pp. 327–337, 2009.

51. D. Rose, W. Thomas, and C. Holm, “Segregation of recombined chromosomes in meiosis i requires dna topoisomerase ii,” Cell, vol. 60, no. 6, pp. 1009–1017, 1990.

52. C. Holm, T. Goto, J. C. Wang, and D. Botstein, “Dna topoisomerase ii is required at the time of mitosis in yeast,” Cell, vol. 41, no. 2, pp. 553–563, 1985.

53. H. Técher, S. Koundrioukoff, A. Nicolas, and M. Debatisse, “The impact of replication stress on replication dynamics and dna damage in vertebrate cells,” Nature Reviews Genetics, vol. 18, no. 9, pp. 535–550, 2017.

54. M. K. Zeman and K. A. Cimprich, “Causes and consequences of replication stress,” Nature cell biology, vol. 16, no. 1, pp. 2–9, 2014.

55. D. Hanahan and R. A. Weinberg, “Hallmarks of cancer: the next generation,” cell, vol. 144, no. 5, pp. 646–674, 2011.

56. M. Macheret and T. D. Halazonetis, “Dna replication stress as a hallmark of cancer,” Annual Review of Pathology: Mechanisms of Disease, vol. 10, no. 1, pp. 425–448, 2015.

57. J. Ray, P. R. Munn, A. Vihervaara, J. J. Lewis, A. Ozer, C. G. Danko, and J. T. Lis, “Chromatin conformation remains stable upon extensive transcriptional changes driven by heat shock,” Proceedings of the National Academy of Sciences, vol. 116, no. 39, pp. 19431–19439, 2019.

58. L. Li, X. Lyu, C. Hou, N. Takenaka, H. Q. Nguyen, C.-T. Ong, C. Cubeñas-Potts, M. Hu, E. P. Lei, G. Bosco, et al., “Widespread rearrangement of 3d chromatin organization underlies polycomb-mediated stressinduced silencing,” Molecular cell, vol. 58, no. 2, pp. 216–231, 2015.

59. S. Korsak and D. Plewczynski, “Loopsage: An energy-based monte carlo approach for the loop extrusion modelling of chromatin,” Methods, 2024.

60. M. G. Gauthier, P. Norio, and J. Bechhoefer, “Modeling inhomogeneous dna replication kinetics,” PLoS one, vol. 7, no. 3, p. e32053, 2012.

61. A. P. de Moura, R. Retkute, M. Hawkins, and C. A. Nieduszynski, “Mathematical modelling of whole chromosome replication,” Nucleic acids research, vol. 38, no. 17, pp. 5623–5633, 2010.

62. R. Yousefi and M. Rowicka, “Stochasticity of replication forks’ speeds plays a key role in the dynamics of dna replication,” PLoS computational biology, vol. 15, no. 12, p. e1007519, 2019.

63. J. Bechhoefer and N. Rhind, “Replication timing and its emergence from stochastic processes,” Trends in Genetics, vol. 28, no. 8, pp. 374–381, 2012.

64. T. Gross and B. Blasius, “Adaptive coevolutionary networks: a review,” Journal of the Royal Society Interface, vol. 5, no. 20, pp. 259–271, 2008.

65. J. Toruniewska, K. Suchecki, and J. A. Holyst, “Unstable network fragmentation in co-evolution of potts spins and system topology,” Physica A: Statistical Mechanics and its Applications, vol. 460, pp. 1–15, 2016.

66. P. Fronczak, A. Fronczak, and J. A. Holyst, “Self-organized criticality and coevolution of network structure and dynamics,” Physical Review E—Statistical, Nonlinear, and Soft Matter Physics, vol. 73, no. 4, p. 046117, 2006.

67. M. V. Imakaev, G. Fudenberg, and L. A. Mirny, “Modeling chromosomes: Beyond pretty pictures,” FEBS letters, vol. 589, no. 20, pp. 3031–3036, 2015.

68. T. Yang, F. Zhang, G. G. Yardimci, F. Song, R. C. Hardison, W. S. Noble, F. Yue, and Q. Li, “Hicrep: assessing the reproducibility of hi-c data using a stratum-adjusted correlation coefficient,” Genome research, vol. 27, no. 11, pp. 1939–1949, 2017.

69. O. Ursu, N. Boley, M. Taranova, Y. R. Wang, G. G. Yardimci, W. Stafford Noble, and A. Kundaje, “Genomedisco: a concordance score for chromosome conformation capture experiments using random walks on contact map graphs,” Bioinformatics, vol. 34, no. 16, pp. 2701–2707, 2018.

70. J. Kubica, S. Korsak, A. B. Clerkin, D. Kouril, D. Schirman, D. Yadavalli, K. Banecki, M. Kadlof, B. Busby, and D. Plewczynski, “The challenge of chromatin model comparison and validation a project from the first international 4d nucleome hackathon,” bioRxiv, pp. 2024–10, 2024.

71. K. Banecki, S. Korsak, and D. Plewczynski, “Advancements and future directions in single-cell hi-c based 3d chromatin modeling,” Computational and Structural Biotechnology Journal, 2024.

72. E. Lieberman-Aiden, N. L. Van Berkum, L. Williams, M. Imakaev, T. Ragoczy, A. Telling, I. Amit, B. R. Lajoie, P. J. Sabo, M. O. Dorschner, et al., “Comprehensive mapping of long-range interactions reveals folding principles of the human genome,” science, vol. 326, no. 5950, pp. 289–293, 2009.

73. G. Le Treut, F. Képès, and H. Orland, “A polymer model for the quantitative reconstruction of chromosome architecture from hic and gam data,” Biophysical journal, vol. 115, no. 12, pp. 2286–2294, 2018.

74. S. Shinkai, M. Nakagawa, T. Sugawara, Y. Togashi, H. Ochiai, R. Nakato, Y. Taniguchi, and S. Onami, “Phi-c: deciphering hi-c data into polymer dynamics,” NAR genomics and bioinformatics, vol. 2, no. 2, p. lqaa020, 2020.

75. S. Shinkai, H. Itoga, K. Kyoda, and S. Onami, “Phi-c2: interpreting hi-c data as the dynamic 3d genome state,” Bioinformatics, vol. 38, no. 21, pp. 4984–4986, 2022.

76. C. Zou, Y. Zhang, and Z. Ouyang, “Hsa: integrating multitrack hi-c data for genome-scale reconstruction of 3d chromatin structure,” Genome biology, vol. 17, pp. 1–14, 2016.

77. O. Oluwadare, M. Highsmith, and J. Cheng, “An overview of methods for reconstructing 3-d chromosome and genome structures from hi-c data,” Biological procedures online, vol. 21, pp. 1–20, 2019.

78. L. Fiorillo, S. Bianco, A. M. Chiariello, M. Barbieri Esposito, C. Annunziatella, M. Conte, A. Corrado, A. Prisco, A. Pombo, et al., “Inference of chromosome 3d structures from gam data by a physics computational approach,” Methods, vol. 181, pp. 70–79, 2020.

79. M. Di Stefano, J. Paulsen, T. G. Lien, E. Hovig, and C. Micheletti, “Hi-c-constrained physical models of human chromosomes recover functionally-related properties of genome organization,” Scientific reports, vol. 6, no. 1, p. 35985, 2016.

80. D. Bau and M. A. Marti-Renom, “Genome structure determination via 3c-based data integration by the integrative modeling platform,” Methods, vol. 58, no. 3, pp. 300–306, 2012.

81. G. Shi and D. Thirumalai, “A maximum-entropy model to predict 3d structural ensembles of chromatin from pairwise distances with applications to interphase chromosomes and structural variants,” Nature Communications, vol. 14, no. 1, p. 1150, 2023.

82. H. Chai, X. Huang, G. Xiong, J. Huang, K. K. Pels, L. Meng, J. Han, D. Tang, G. Pan, L. Deng, et al., “Tri-omic single-cell mapping of the 3d epigenome and transcriptome in whole mouse brains throughout the lifespan,” Nature Methods, pp. 1–14, 2025.

83. T. J. Stevens, D. Lando, S. Basu, L. P. Atkinson, Y. Cao, and F. Steven, “3d structure of individual mammalian genomes studied by single cell hi-c,” vol. 544, pp. 59–64, 2017.

84. L. Affonso, R. Bissacot, E. O. Endo, and S. Handa, “Longrange ising models: Contours, phase transitions and decaying fields,” Journal of the European Mathematical Society, 2024.

85. J.-C. Anglès d’Auriac and F. Iglói, “Phase transitions of the random-bond potts chain with long-range interactions,” Physical Review E, vol. 94, no. 6, p. 062126, 2016.

86. I. Lecce, M. Picco, and R. Santachiara, “Magnetic exponent for the long-range bond-disordered potts model,” Physical Review E, vol. 110, no. 6, p. 064154, 2024.

87. E. Bayong, H. Diep, and V. Dotsenko, “Potts model with longrange interactions in one dimension,” Physical review letters, vol. 83, no. 1, p. 14, 1999.

88. M. Biskup, L. Chayes, and N. Crawford, “Mean-field driven first-order phase transitions in systems with long-range interactions,” Journal of Statistical Physics, vol. 122, pp. 1139–1193, 2006.

89. W. Schwarzer, N. Abdennur, A. Goloborodko, A. Pekowska, G. Fudenberg, Y. Loe-Mie, N. A. Fonseca, W. Huber, C. H. Haering, L. Mirny, et al., “Two independent modes of chromatin organization revealed by cohesin removal,” Nature, vol. 551, no. 7678, pp. 51–56, 2017.

90. J. Nuebler, G. Fudenberg, M. Imakaev, N. Abdennur, and L. Mirny, “Chromatin organization by an interplay of loop extrusion and compartmental segregation,” Biophysical Journal, vol. 114, no. 3, p. 30a, 2018.

91. A. Goloborodko, J. F. Marko, and L. A. Mirny, “Chromosome compaction by active loop extrusion,” Biophysical journal, vol. 110, no. 10, pp. 2162–2168, 2016.

92. S. C.-H. Yang, M. G. Gauthier, and J. Bechhoefer, “Computational methods to study kinetics of dna replication,” DNA Replication: Methods and Protocols, pp. 555–573, 2009.

93. A. Baker and J. Bechhoefer, “Inferring the spatiotemporal dna replication program from noisy data,” Physical Review E, vol. 89, no. 3, p. 032703, 2014.

94. R. Retkute, C. A. Nieduszynski, and A. De Moura, “Mathematical modeling of genome replication,” Physical Review E—Statistical, Nonlinear, and Soft Matter Physics, vol. 86, no. 3, p. 031916, 2012.

95. H. Zhang and J. Bechhoefer, “Reconstructing dna replication kinetics from small dna fragments,” Physical Review E—Statistical, Nonlinear, and Soft Matter Physics, vol. 73, no. 5, p. 051903, 2006.

96. D. Löb, N. Lengert, V. Chagin, M. Reinhart, C. Casas-Delucchi, M. Cardoso, and B. Drossel, “3d replicon distributions arise from stochastic initiation and domino-like dna replication progression,” Nature communications, vol. 7, no. 1, p. 11207, 2016.

97. J. H. Yang, H. B. Brandão, and A. S. Hansen, “Dna double-strand break end synapsis by dna loop extrusion,” Nature communications, vol. 14, no. 1, p. 1913, 2023.

98. D. D’Asaro, M. M. Tortora, C. Vaillant, J.-M. Arbona, and D. Jost, “Dna replication and polymer chain duplication reshape the genome in space and time,” Physical Review X, vol. 14, no. 4, p. 041020, 2024.

99. D. D’Asaro, J.-M. Arbona, V. Piveteau, A. Piazza, C. Vaillant, and D. Jost, “Genome-wide modeling of dna replication in space and time confirms the emergence of replication specific patterns in vivo in eukaryotes,” bioRxiv, pp. 2025–04, 2025.

100. T. Nagano, Y. Lubling, T. J. Stevens, S. Schoenfelder, E. Yaffe, W. Dean, E. D. Laue, A. Tanay, and P. Fraser, “Single-cell hi-c reveals cell-to-cell variability in chromosome structure,” Nature, vol. 502, no. 7469, pp. 59–64, 2013.

101. D. J. Massey and A. Koren, “High-throughput analysis of single human cells reveals the complex nature of dna replication timing control,” Nature Communications, vol. 13, no. 1, p. 2402, 2022.

102. P. Eastman, J. Swails, J. D. Chodera, R. T. McGibbon, Y. Zhao, K. A. Beauchamp, L.-P. Wang, A. C. Simmonett, M. P. Harrigan, C. D. Stern, et al., “Openmm 7: Rapid development of high performance algorithms for molecular dynamics,” PLoS computational biology, vol. 13, no. 7, p. e1005659, 2017.

103. P. Eastman, R. Galvelis, R. P. Peláez, C. R. Abreu, S. E. Farr, E. Gallicchio, A. Gorenko, M. M. Henry, F. Hu, J. Huang, et al., “Openmm 8: molecular dynamics simulation with machine learning potentials,” The Journal of Physical Chemistry B, vol. 128, no. 1, pp. 109–116, 2023.

104. J. M. Finn, Classical mechanics. Jones & Bartlett Publishers, 2009.

105. M. E. Newman and G. T. Barkema, Monte Carlo methods in statistical physics. Clarendon Press, 1999.

106. M. Kardar, Statistical physics of particles. Cambridge University Press, 2007.

107. D. Jost, P. Carrivain, G. Cavalli, and C. Vaillant, “Modeling epigenome folding: formation and dynamics of topologically associated chromatin domains,” Nucleic acids research, vol. 42, no. 15, pp. 9553–9561, 2014.

108. F. S. Bates and G. H. Fredrickson, “Block copolymer thermodynamics: theory and experiment,” Annual review of physical chemistry, vol. 41, no. 1, pp. 525–557, 1990.

109. R. Zhou and Y. Q. Gao, “Polymer models for the mechanisms of chromatin 3d folding: review and perspective,” Physical Chemistry Chemical Physics, vol. 22, no. 36, pp. 20189–20201, 2020.

110. A. Oji, L. Choubani, H. Miura, and I. Hiratani, “Structure and dynamics of nuclear a/b compartments and subcompartments,” Current Opinion in Cell Biology, vol. 90, p. 102406, 2024.

111. K. Samejima, J. H. Gibcus, S. Abraham, F. Cisneros-Soberanis, I. Samejima, A. J. Beckett, N. Puaceková, M. A. Abad, C. Spanos, B. Medina-Pritchard, et al., “Rules of engagement for condensins and cohesins guide mitotic chromosome formation,” Science, vol. 388, no. 6743, p. eadq1709, 2025.

112. S. K. Lam, A. Pitrou, and S. Seibert, “Numba: A llvm-based python jit compiler,” in Proceedings of the Second Workshop on the LLVM Compiler Infrastructure in HPC, pp. 1–6, 2015.

